# Paw posture is a robust indicator for injury, pain, and age

**DOI:** 10.1101/2025.11.13.688005

**Authors:** Christian O. Pritz, Sofia Dotta, Philip A. Freund, Giada Musso, Michael Bader, Nataliya Okladnikov, Franziska Rother, Letizia Marvaldi

**Affiliations:** Department of Neuroscience "Rita Levi Montalcini", Neuroscience Institute Cavalieri Ottolenghi, University of Turin, 10043 Orbassano (TO), Italy; Max Delbrück Center for Molecular Medicine, 13125 Berlin, Germany; Center for Structural and Cellular Biology in Medicine, Institute of Biology, University of Lübeck, 23538 Lübeck, Germany; Charité - Universitätsmedizin Berlin, 10117 Berlin, Germany; Department of Biomolecular Sciences, Weizmann Institute of Science, Rehovot 76100, Israel

**Author notes:** Last authors (Franziska Rother, Letizia Marvaldi). equally contributed (Sofia Dotta, Philip A. Freund, Giada Musso).

## Abstract

Inferring biological states from animal behavior is a crucial but challenging step in biomedical discovery that is constrained by variability and labour-intensive assays, even with AI-powered tools. Here, we show that simple images of static paws, analyzed by our custom keypoint segmentation AI-tool provide accurate read-outs for a wide array of physiological states. Without invasive testing, our method detects postural changes associated with nerve injury, acute pain, aging, and the genetic loss of kpna4, a regulator of paw innervation. Leveraging the toe-spread-reflex, a spinal-circuit driven response, the approach requires no habituation and shows low behavioral variability. Individual digits emerge as biomarkers for internal states with digit V indicating neuropathic pain during nerve damage and digit I reflecting loss of kpna4. Our model is freely available and can readily be adapted to other tasks or species. These findings establish unstimulated paw posture as a scaleable, low-cost, readout for biological states.

## Introduction

Estimating internal physiological changes of an animal from behavior is a very challenging task. Yet, it is important and often the first step in biomedical scientific discovery. Fortunately, the scientific community can rely on many established behavioral assays to mine for changes in precisely defined territories within the animal’s neurophysiology, general physiology, and genetics. Despite these robust methods, converting behavioral observations to reliable metrics and detecting significant and biologically relevant changes is limited by high variability, and often low effect size. Additionally, many of these assays require skilled and intensive animal handling that reduces the size available to offset inherent variability. In the past decade, progress in artificial intelligence (AI) and the rise of animal pose segmentation models^1–5^ lead to an unprecedented increase in the amount of data that can be collected. These deep-learning-powered tools can help eliminate human bias in the scoring of animal behavior and quantifying spontaneous behavior^6–9^. However, increased data volume and complexity of the data requires more analytical effort and more statistical expertise to accurately link behavioral variation to relevant biological phenotypes^10^; accurate behavioral analysis still remains a challenge.

Assessing the degree of peripheral nerve damage via behavioral assays, for instance, is particularly demanding. It requires a surgical intervention to induce a precise nerve injury, which in turn can lead to unwanted behavioral variability. Then, the effects of and recovery from injury are usually estimated from a set of non-trivial assays. Some tests probe sensory function, including the Hargreaves and von Frey test. In the Hargreaves test, the mouse’s paw is exposed to a 52-58 degree-heated metal probe to estimate pain threshold by the latency to paw withdrawal^11^. In the von Frey test, nylon filaments are pushed against the ventral side of the hind paw, measuring the force threshold needed to elicit paw withdrawal^12^. Both tests are robust but require habituation and precise manual handling of the animal. Motor performance can also be used as a proxy for nerve recovery. Two widely employed methods are the CatWalk test^13^, which quantifies gait by recording the mouse’s walking pattern and the rotarod test^14^ which assesses motor coordination and balance by the mouse’s ability to remain on a rotating rod. Both assays demand active engagement in the task and can be influenced by learning effects, which may substantially confound the results. These experimental procedures inevitably cause discomfort to the animal and are prone to noise unrelated to the physiological process of nerve recovery.

In our previous efforts, we identified importin alpha 3 (kpna4) as a regulator of chronic pain during the late phase of recovery from spared nerve injury (SNI) using the aforementioned methods. During extensive behavioural testing^15^, we observed a striking paw posture that is indicative of severe nerve damage and can be confidently used to classify the injury state. Inspired by this observation and preceding successful uses of AI to study pain and behavior^16,17^, we custom-trained Meta’s detectron2 suite^18^ to identify and quantify paw posture and to infer virtual paw skeletons from images of static paws thereby reducing experimental effort. These images can be acquired during routine animal handling using a low-cost approach based on consumer cameras eliminating the need for habituation or a dedicated setup. Using this powerful tool, we show that analysis of the paw posture can unmask a wider array of internal variables of the animal besides injury state during SNI. We demonstrate that the virtual skeleton of a static hind paw can accurately highlight responses to acute pain, reveal the postural signature of aging, and even uncover postural deviations caused by the kpna4 knockout (KO) loss. Our analysis identified intuitive postures that can serve both as biomarkers for visual inspection and targets for quantitative analysis. We observed that the positioning of digit V after SNI reflects the pain sensitivity, aging strongly modulates digit II and V, and kpna4 KO markedly alters positioning of digit I. Our results demonstrate that unevoked limb paw is a simple and robust readout for a variety of physiological parameters.

## Results

### Paw opening angle indicates sciatic nerve injury

As published previously, loss or reduction of kpna4 in mouse dorsal root ganglion neurons results in a better recovery from sciatic nerve injury (SNI) and reduction of neuropathic pain three months after SNI^15^. After performing SNI (**Fig 1 A,B**), we monitored recovery with the von Frey sensory test (**Fig 1C**) and CatWalk for three months. To test whether kpna4 affects recovery, it was depleted in motor and sensory neurons using AAV9-mediated shRNA approach prior to the injury. In another group kpna4 was depleted exclusively in sensory neurons after the injury using the AAV PhP.S virus as a vector to model a therapeutic intervention. Both groups showed a significant reduction in their pain thresholds in the von Frey test (**Fig 1D&E**). Besides the behavioral tests, we took images of the hind paw during behavioral tests three months after SNI (**Fig 1F**). Strikingly, SNI-treated paws displayed a closed posture, while digits of the contralateral paw retained a spread posture (**Fig 1G**). This prompted us to ask if there is a significant difference in the total opening angle (TOA, manually measured between digits I and V, **suppl. Fig 1B**) between injured and non-injured paws. As the difference was significant (p=2.24x10^-15^, Watson Williams test, **Fig1 H**), we tested if the differences are robust enough to classify injury status of a paw based on the opening angle. For that purpose, we pooled all data from all experimental groups. Using logistic regression on the toe opening angle, we accurately classified injured and non-injured paws with an accuracy and a F1 score of 0.83 and 0.83, respectively (**Fig 1IJ**). This observation indicates that the paw posture is indicative of the nerve injury status. However, when examining the overlap in toe spreading angle (**Fig 1I**), the imperfect separation of groups is evident. Uninjured paws might spontaneously close and injured paws might open due to varying severity of injury or partial recovery in some of the virus-treated experimental groups. These observations highlighted that injury significantly alters paw posture, but also encouraged us to pursue a more automated measurement of the geometry of the paw posture to gain more robust insights.

**Figure 1.**
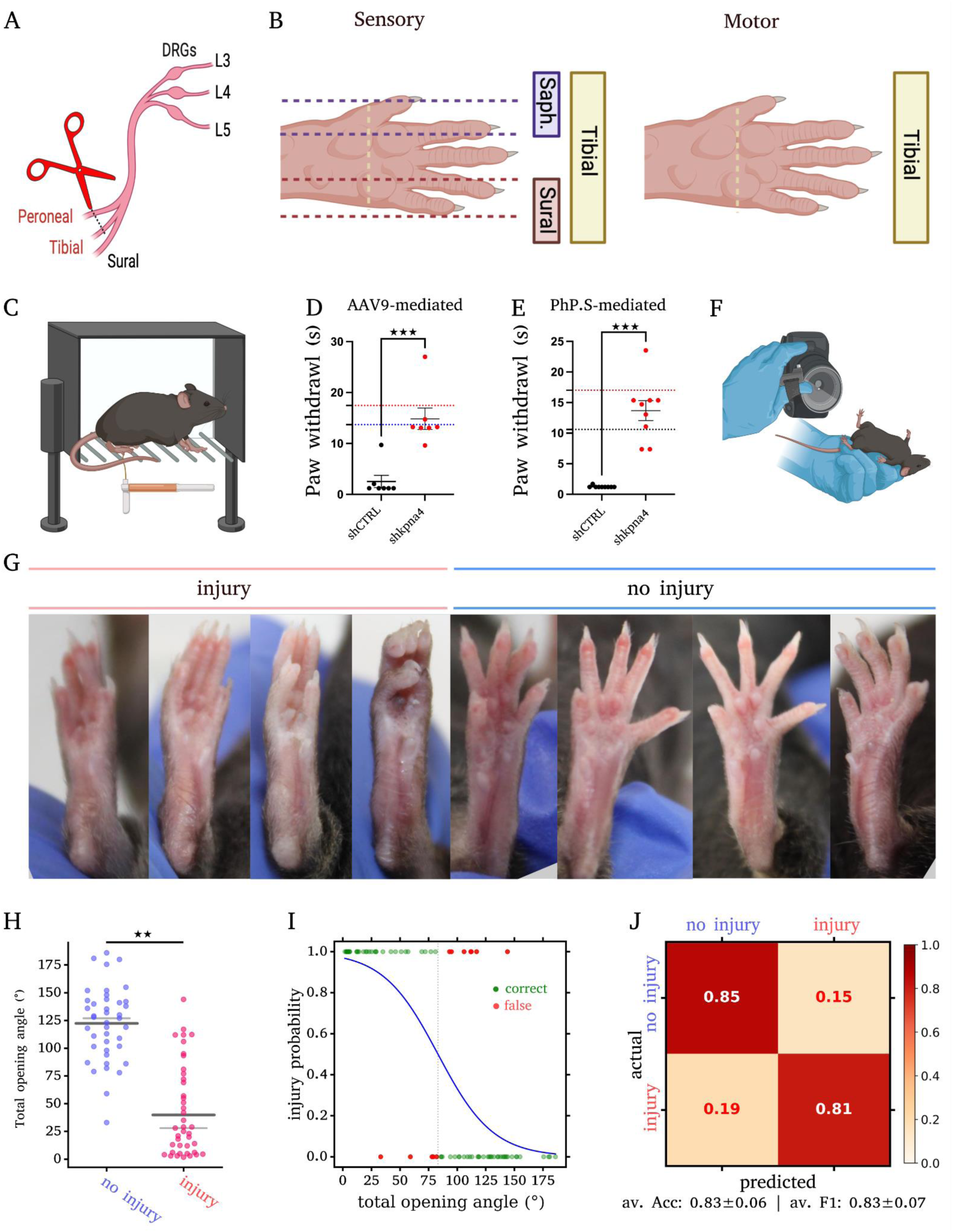
Paw posture is indicative of injury state. **A** Spared nerve injury model: the peroneal and tibial were cut and ligated, the sural nerve remained intact. **B** Neural territories for hindpaw innervation for sensory and motor innervation **C** Von Frey hair filament test was performed in the external region of the hindpaw, where the sural nerve is innervating. **D-E** graphs adjusted from Marvaldi et al., 2020 both AAV9 (sensory and motor neurons, **D**) and PhP.S-mediated (sensory-specific, **E**) shRNA-based kpna4 depletion leads to recovery of mechanical threshold measured in mN (black dashed line, contralateral paw WT mice; red dashed line, contralateral paw KO mice). **G** Examples of injured and non-injured paws. Note the closed posture in the injured paw. **H** When measuring and plotting the angle between digit I and V, a significant difference between injured and non-injured paws can be appreciated (Watson Williams test, p = 2.24 x 10^-15^). The angle can also be used to accurately classify the injury status by logistic regression (**I**) yielding sufficient classification accuracy. **J**, confusion matrix from I denoting relative counts per class for both the actual and the predicted labels. Av. Acc. = Average accuracy; av. F1 = average binary F1 score; n = 41 animals; * p<0.05, ** p<0.01, *** p<0.001

### Keypoint segmentation accurately infers paw geometry

To rapidly mine a virtual skeleton from paw photographs, we used a mask R-CNN model from Meta’s detectron2 suite, a powerful tool to extract the paw position and keypoints defining the paw posture. Therefore, we retrained one of Meta’s detectron2 keypoint R-CNN models to recognize 15 points defining the virtual skeleton of the paw (**Fig 2A**). Based upon previously compiled images^15^ and images from routine care, we created an annotated training set consisting of 959 images. These are photos from mice held in the neck-fold grip taken with either consumer SLRs or mobilephone cameras. Images showed close-ups of hind paws. Keypoints and bounding boxes were then annotated by three expert annotators. In our dataset, paws displayed three prominent poses: an open posture with toes spread out in varying degrees, a closed posture in which most digits would align, and a clenched posture. While the first posture was more frequent, the latter two were rarer and more associated with the injured paws. To address underperformance on these rare postures, we overrepresented clenched and closed postures when augmenting the dataset with rotations, scaling, translation, colour space permutations, and composite image creation (**suppl. Fig. 1**). The model was trained for 319k iterations or 34 epochs. Evaluating the predictions of our trained model on the evaluation set reveals an average precision of 0.97 for keypoints (**Fig 2C**) on our held-out test set. Calculating the mean squared error (MSE) for deviations in the keypoint placement and resulting angular deviations (**Fig 2D-H**) unmasked several properties that influenced the overall performance of our model. Strikingly, postures that are self-occluding (**Fig 2B**) accumulate more error in keypoint placement (**Fig 2D&E**). To quantify self-occlusion in paws, we summed the number of keypoints annotated as non-visible per paw: no self-occlusion tolerates up to one invisible, moderate includes 2 or 3 invisible, and severe when more than 3 keypoints are not visible. Severely self-occluded postures were also difficult to annotate for human scorers to annotate and required consensus annotation. Consequently, it is not surprising that these postures are more error prone. Since convolutional neural networks attend to texture ^19,20^, we correlated the sharpness of the paw (expressed by the Laplacian variance) and the MSE, denoting more MSE in poorly focused images, which can be a limiting factor when acquiring images for analysis. We checked if paw size relative to the image (the paw resolution) influences the error, and found that higher MSE is associated with smaller paws (**Fig 2H**). We next asked how these placement errors would propagate into metrics like the TOA (**Fig 1**). Therefore, we calculated the average deviation for each of 153 pairwise angles between the edges and found an overall 7.4-degree deviation and 9.5 degrees for the TOA. Interestingly, a low number of large placement errors (residuals > 3 σ, arrowhead in **Fig 2H**) have disproportionate influence, suggesting that low number of catastrophic misplacements deteriorate the overall accuracy in keypoint placement. Lastly, projecting the placement error back onto the paw (**Fig 2J**) reveals that the tips of the digits are associated with relatively low placement error, while the bases and joints of the digits II-IV are associated with higher error rate, presumably due to more frequent occlusion.

**Figure 2.**
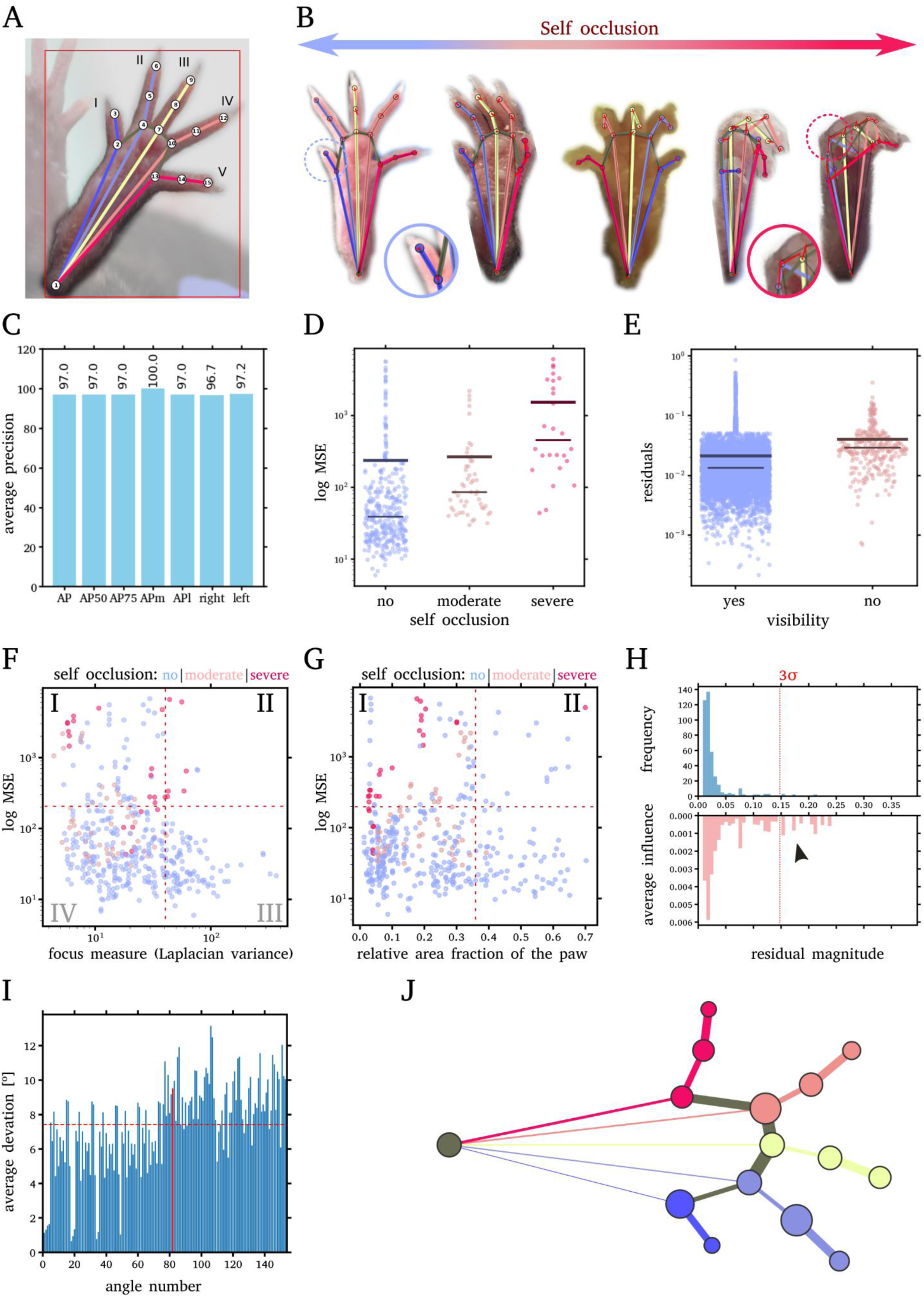
Model performance on the held out test dataset. **A** 15 keypoints were used to train the model. **B** Paw postures can be self-occluding when paws are clenched or turned (coloured lines indicate model predictions, solid black lines ground truth, red arrows indicate errors and E). Note that self-occluded postures incur larger errors, see insets. **C** Average precision over multiple thresholds of object keypoint similarity (75%, 50% medium and large objects, and averages for left and right paws separately) on test instances. **D** Mean squared error (MSE) of individual paws with differing degrees of self-occlusion. **E** placement error (residuals) for individual keypoints; note that keypoints that are occluded are associated with higher error **F** Images of less focused paws (lower Laplacian variance), incur higher MSE (compare sectors I and II). **G** Paws with lower resolution (lower area fraction in the image) are more likely to have higher error (compare sectors I and II). **H** Residual diagnostics: Upper histogram denotes magnitude of residuals, lower histogram its relative impact on the mean. Note that a low number of large errors (> 3 σ, see arrowhead) are disproportionately affecting the mean. **I** Average deviation of all pairwise angles between edges. The dotted line denotes overall mean and the red bar denotes the error of the TOA. Note that from angle 75, pairwise angles involve digit segments and thus show more variability because of higher degrees of freedom **J** Placement error of keypoints (circle radius is 90 % error probable) and the relative contribution of each edge to angular errors (width of edges). Endpoints of digits, digits I, II, V show relatively low error, while the error of the remaining keypoints is increased by self-occlusion in certain paw postures.

These results show our model performs best on sharply focused images of single paws with good resolution. To mitigate inaccuracy by keypoint wobble (especially in self-occluding poses that strongly deteriorate the overall accuracy), we implement a user interface that allows the user to correct eventual errors by simply dragging the predicted keypoints. In the following, data presented was processed using this semi-automatic pipeline.

Once trained, we applied our model to all virtual skeletons in this study (see **suppl Fig 3**) to quantify the postures. The keypoint coordinates formed the basis for quantitative analysis of paw geometry yielding a variety of metrics (see **suppl Fig 1**). We then subjected the resulting virtual skeletons to a principal component analysis (PCA) capturing the main axes of postural variability. User-annotated postures occupied distinct territories in principal component (PC) space (**suppl Fig 3A-D**). All subsequent classification tasks used PC scores from this global PCA - a strategy that lowers accuracy but reduces the risk of overfitting. Subjecting the PC scores that separated the postures (1,3,6,8,10, and 17) to a random forest classifier identified the three intuitive, user-annotated postures with good accuracy (av. F1 score 0.83±0.04, **suppl. Fig 3E**). Hierarchical clustering of the same PCs groups most ‘clenched’ and ‘closed’ postures together with some posturally similar ‘open’ paws, while the remaining clusters are predominantly composed of ‘open’ (**suppl Fig 3F**). An elbow-plot of within-cluster variance shows no distinct inflection point indicating an optimal number of clusters (**suppl Fig 3G**). Because closing and clenching are fluid motion sequences with many transitional postures that would be hard to delineate, we restricted downstream analysis to the three intuitive postures.

### Paw posture indicates injury history

With the virtual skeleton of the paw accessible, we could infer injury history from the geometry of the virtual skeleton. When performing PCA on the keypoints injured and non-injured paws separated well in PC space (**Fig 3A**). This separation accurately distinguished between injured and non-injured paws (av. F1 score 0.90±0.03, Fig 3C) using a random forest classifier based on PCs in which we find injury-related separation (1,3,8,11,12,13, and 19). Calculating the average virtual skeletons of the injured and non-injured groups from the centers-of-mass of the PCs showed a ‘closed’ and ‘open’ conformation, respectively (**Fig 3B**), which is consistent with user annotation of postures. The ‘closed’ and ‘clenched’ configurations were significantly enriched in the injured group (**Fig 3E**, Fisher’s exact test) supporting our initial intuition (**Fig 1G**). Together, this suggests the closed paw posture is indicative of past SNI.

**Figure 3.**
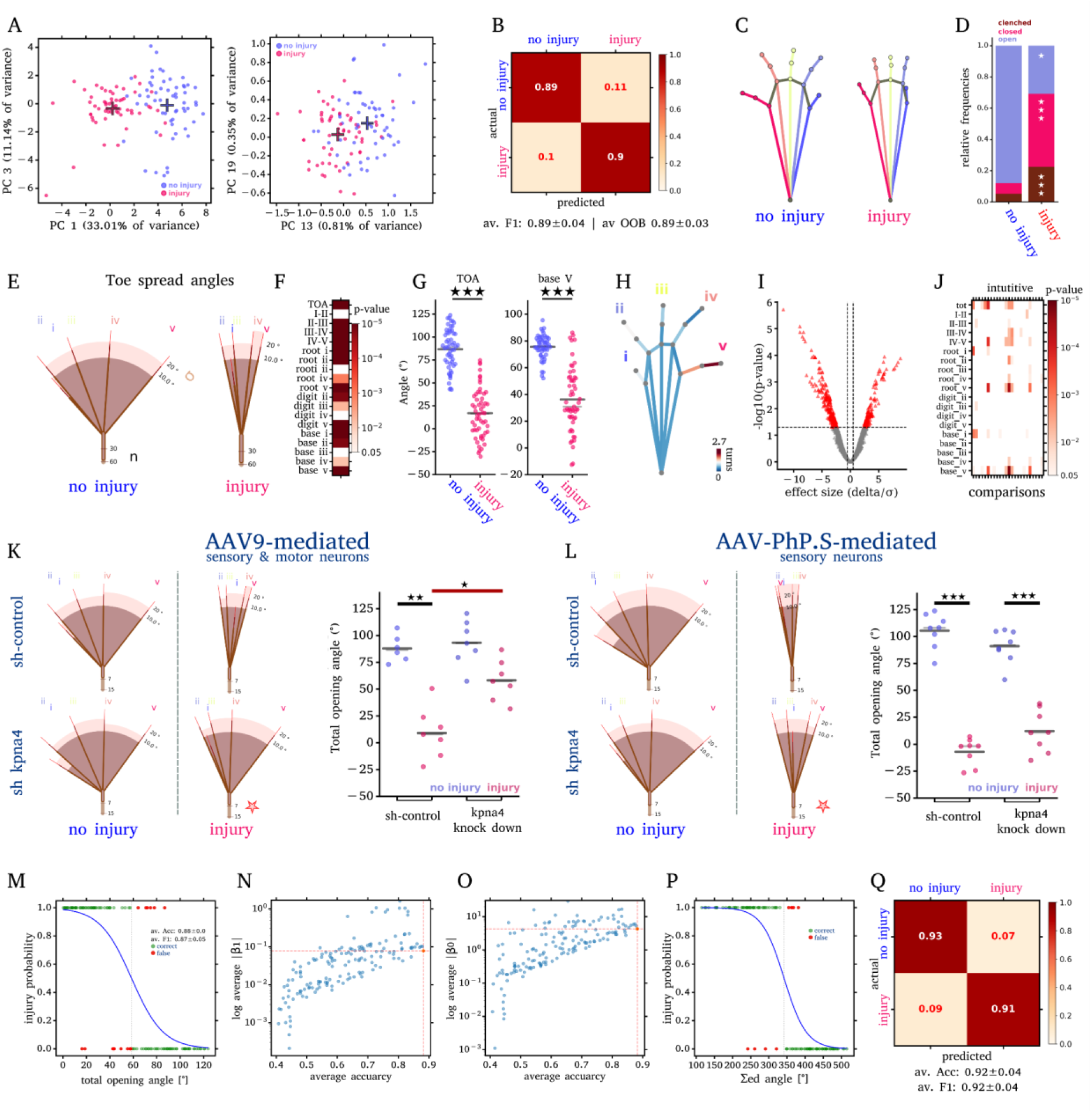
Closed paw posture indicates injury. **A** PCA of keypoints shows distinct clusters for injured and non-injured paws. **B** Confusion matrix from random-forest (RF, 500 trees) classifier injured vs non-injured paws based on PC scores (1,3,8,11,12,13, and 19). **C** Group-specific postures from the centre-of-mass (crosses in A) reveals a ‘closed’ posture in injured groups. **D** Relative frequency of postures for injured and non-injured paws. Note that relative frequencies of postures changed significantly (Fisher’s exact test, Bonferroni corrected) **E** The more compact posture of toes of injured paws is evident in a fan plot (see **suppl.** Fig. 1A). **F** Many significant differences exist between injured and non-injured paws (Watson Williams test, FDR corrected, 153 comparisons, p-values color-coded for intuitive angles, see suppl Fig 1B). **G** TOA and base V angle scatter plots. **H** Projecting significant angles back onto the edges, reveals contributions of digits I,II,IV, and in especially digit V. **I-J** Considering treatment-groups in hypothesis testing yields many significant differences (I, volcano plot, J heatmap, 612, comparisons, WW test). **K** AAV9-mediated knock down of kpna4 in motor and sensory neurons leads to significantly larger opening angle after injury (see fan and scatter plots). Note the differences between injured paws treated with sh control and the kpna4 knockdown (red bar). **L** When only sensory neurons are targeted no increase in the treated group is observed. While there is a significant difference in opening angles (red asterisks, J & K) in opening angles between injured paws that were treated with AAV9 and AAV PhP.s. **M** Logistic regression on TOA with 20-fold cross validation. No other angle in the paw geometry yields better accuracy, and it shows a numerically high β_1_ (steep probability transition, **N**) but a small β_0_ (early shift in the probability transition point**, O**). Only a combination of angles yields better accuracy (**P, Q**). Note the better separation and steeper probability transition. * p<0.05, ** p<0.01, *** p<0.001 (Watson Williams tests, FDR corrected) n = 58 animals, A-H & N-R

Next, we analysed the angles between individual edges of the virtual skeleton. The drastically altered geometries in injured paws are evident when plotting the angles between individual toes as a fan plot (**Fig 3E**). When testing for significant differences between all pairwise angles (153 Watson Williams tests, FDR corrected) between injured and control paws pooling all the groups, it is apparent that many angles significantly changed (**Fig 3F&G, suppl Fig 3H&I**). Notably, the TOA showed a significant delta even though the injured group included animals that partially recovered (see **Fig 1D & E**). The significant angular change can be mapped back onto the edges of the virtual skeleton by color-coding the sum of significant angular deltas each altered edge produced (**Fig 3H, suppl. Fig 1C**). Consistent with visual observation (**Fig 3H**, **Fig 1G**), digit I, II and V yielded most of the significant change. Strikingly digit V seemed to contribute disproportionally to the overall postural change. In conclusion, injury caused a lasting change in paw posture driving the paw towards a closed confirmation that is detectable visually and quantifiable.

### Toe motility is partially regained when kpna4 is depleted in motor neurons

Informed by this analysis, we broke down the data into different virus-treated groups to check whether different treatments varied paw posture. Briefly, kpna4 was knocked down in dorsal root ganglion neurons using shRNA targeting either by AAV9 virus (motor and sensory neuron) or AAV-PhP.S (sensory neurons only)^21,22^ around the time point of injury (see material and methods). When accounting for all different subgroups (kpna4 shRNA, controls, different viruses), most of those differences are still detectable (**Fig 3I&J, and suppl Fig. 3 L**).

Comparing the individual toe angles (fan plots) and TOA (scatter plots) in **Fig 3K** and **L**, it is apparent that in all groups the injury significantly reduced the opening of the paw digits. However, there was a significant recovery of TOA specifically in injured paws that also underwent AAV9-mediated kpna4 knockdown but not paws that received AAV-PhP.S-mediated knockdown (compare scatter plots in **Fig 3 K and L**). A better recovery in paw posture is also evident from group-specific postures (**Fig 3J & suppl Fig 3L**). Non-injured paws and paws that underwent SNI injury together with AAV9-mediated kpna4-depletion showed an open posture (**suppl Fig 3J**), while injured paws that received sham treatment (sh control) or AAV-PhP.S mediated knockdown exhibit injury specific postures (compare arrowheads in **suppl Fig. 3J**) This suggests a partial rescue of the phenotype when motor and sensory neurons are targeted, while targeting only sensory neurons after injury is insufficient. Together, these observations would suggest that kpna4 loss in motor and sensory neurons can partially restore toe motility after SNI. It also indicates that quantifying the TOA is a sensitive tool to monitor recovery by quantifying toe motility.

### TOA is a robust indicator of SNI

As the keypoint segmentation provides us with all possible angles in the virtual skeleton, we next tried to confirm whether our initial intuition on the TOA is indeed a robust metric to classify injury history. When we reclassified the paws using a logistic regression on the TOA, we found the classification accuracy (av. F1 score 0.81) in good agreement with our manual measurements (compare **Fig 3M**, with **Fig 1I&J**). To verify whether any other angle between any edge of the virtual skeleton yields a better accuracy, we performed logistic regression on all 153 pairwise angles against the injury status. In the logistic function β_1_ controls the steepness of the probability transition and β_0_ shifts its onset. When we plot β_0_ and β_1_ against the classification accuracy for all coefficients (**Fig 3N,O**), it is apparent that the TOA (marked in red) was consistently found amongst three strongest predictors, while also showing a numerically small β_1_ (**Fig 3N)**, yielding a steep probability transition but a low β_0_ shifting the decision threshold towards smaller angles (**Fig 3O**). We then tested whether combining different angles could further improve accuracy. Indeed, when combining the angles between edges depicted in **suppl Fig 3K**, we obtained better discrimination between injured and non-injured paws (av. F1 score of 0.92±0.04, compare **Fig 3M & P,Q**). These predictive angles are associated with positioning of all the digits **(suppl. Fig. 3K),** together describing the paw opening well. While yielding higher accuracy in classification, a combination of angles is not intuitive as a visual marker. These observations validated the TOA as both a robust visual biomarker for injury history and for quantification.

### Paw posture is indicative of acute pain

The strong correlation between paw posture and injury led us to question if the configuration of the paw could also reflect pain. To test this hypothesis without the bias of injured motor fibers, we conducted capsaicin injections of the hind paw. Animals would receive a single injection of vehicle control, 25, or 50 mg capsaicin per kg bodyweight. Paw postures were recorded before injection (10 and 0 min) and 10, 20, 30, 40 seconds immediately after the injection and at 5 and 60 minutes. Leveraging the virtual skeleton, we first analyzed the overall postures of the injected paws. Strikingly, penetration by the needle caused all the paws to clench. After extraction of the needle, paws would progressively change into the open posture. While the opening kinetics of 25 mg.kg^-1^ dosage was similar to the vehicle control, the clenched posture persisted the longest in the 50 mg capsaicin group (**Fig 4A**). Analyzing posture distribution following injection reveals postural variability in all injected groups for the minute following the injection (**Fig 4B** & **suppl Fig 4**), while after 5 minutes postures become more homogenous and comparable. Due to the limited group sizes and conservatism of Fischer’s exact test, we could not detect changes in postural frequencies between vehicle control and capsaicin injected groups. However, it is noteworthy to point out that the ‘closed’ posture was only observed in capsaicin-injected groups (**suppl Fig 5A**, **Fig 4B**). For statistical comparisons of paw geometry, we analyzed the pre-injection, 20 second, 5 min and 60 min time points. We compared the two capsaicin concentrations with each other and with vehicle controls, the vehicle control group to the non-injected side, and successive timepoints within each group. Unsurprisingly, many angles were changed by injections (**Fig. 4D-F)**, while we could detect statistically significant differences even one hour after injection like the overall increased TOA (p<10^-6^, permutation test, 10k-fold, FDR corrected) and increased prominence of digit I from the paws in all injected groups (p=0.012, **Fig 5C**). Due to the high postural variability, individual angle differences are not particularly informative, however, the number of significant differences highlights the overall postural change between two groups (**Fig. 4E**). Only one angle significantly differed between vehicle control and 25 mg group, whereas more than 35 differed between 50 mg group and vehicle controls. Thus 50 mg capsaicin induced robust postural changes beyond the effect of needle injection visible in the vehicle control, while the lower dosage is posturally very similar to the vehicle-control paw. Notably, the largest shift occurred between the vehicle control group and the non-injected contralateral paw underscoring the strong postural effect of the injection itself. All groups exhibited significant postural changes over time when analyzed within groups. The 50 mg group stood out by showing the least number of changes, likely due to its high positional variability immediately after the injection. Projecting these changes back onto the edges of the virtual skeleton (**Fig. 4G**) reveals that injection alone significantly modulates all digits (comparing non-injected vs. vehicle control paws). By contrast, capsaicin-specific postural changes are weaker (**Fig 4G**) but further alter the positioning of digits II, III, IV (capsaicin paws vs. vehicle control). This indicates that the penetration by the needle induces robust postural change in all digits which are further modulated by the effects of capsaicin.

**Figure 4.**
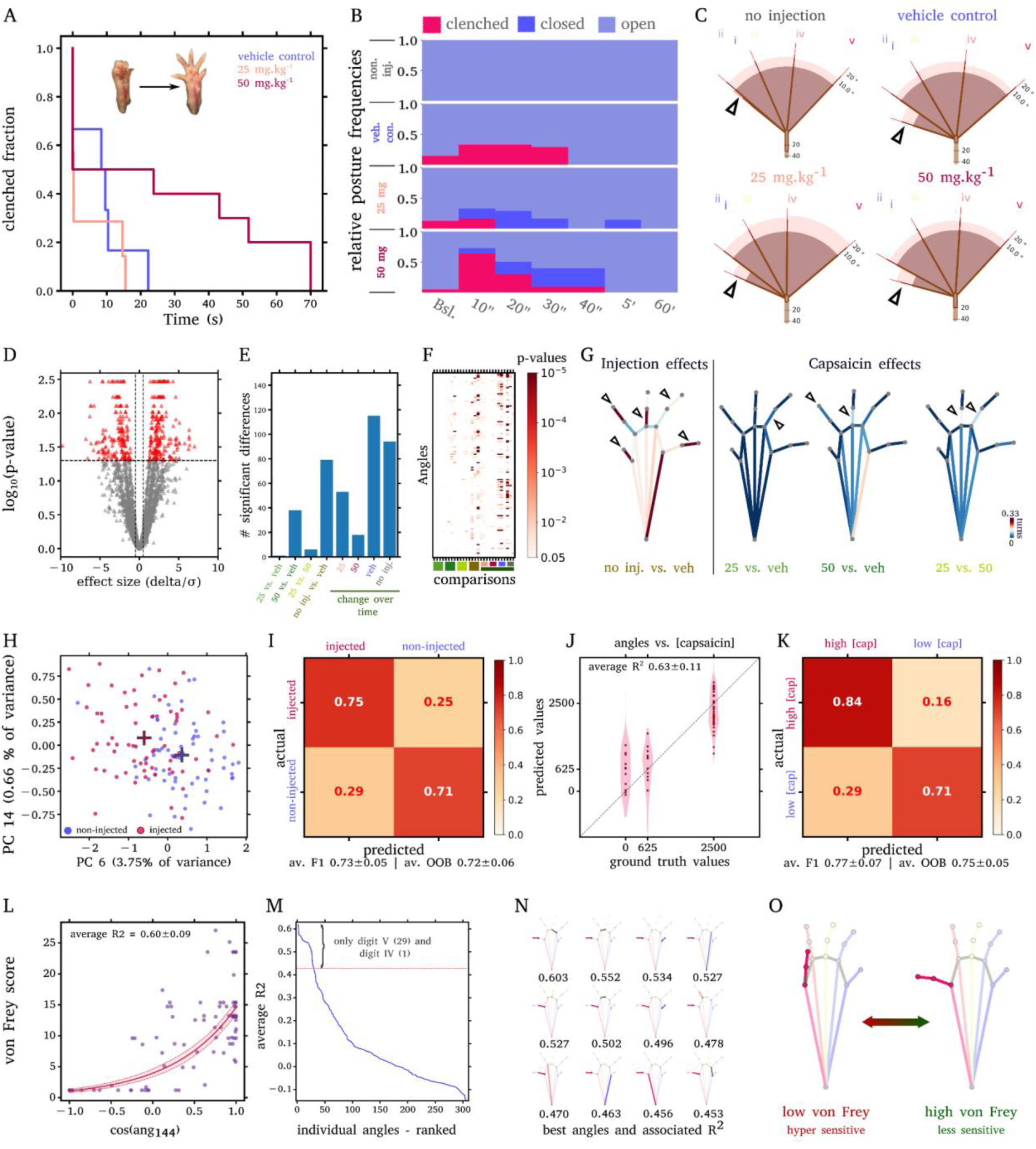
Pain modulates hind paw posture. **A** The fraction of clenched paws over time, **B** relative fraction of paw postures for different time points **C** Inter-digit angels per group one hour after injection. **D** many angles are significantly changed by injection (permutation tests, 4284 comparisons, FDR corrected), however there are more differences between vehicle control and non-injected paws than capsaicin-injected and vehicle control (**E**, differences per comparison category). **F** significant differences per comparison. **G** Paw-mapping of significant changes after one hour for each comparison. (Note that all digits are affected in vehicle control vs. non-injected group). **H** Injected (capsaicin & vehicle control) and non-injected paws diverge in PC space. **I** Confusion matrix from random-forest (RF) classification of non-injected and injected paws based on PC scores (2, 4, 5, 7, 12, 16, 10, 23, and 29). **J** Predicted vs. ground truth values from multivariate regression of paw angles vs capsaicin concentration (dots denote predictions from the full model, violins denote predictions from crossvalidation trials). **K** Confusion matrix of the RF classification of paws receiving high and low capsaicin dosage (low: 25 mg.kg^-1^ pooled with vehicle control). **L** Univariate regression of paw angle 144 (see first in M) vs von Frey score three months after SNI (line denotes prediction from full model, shaded area 90 % confidence interval from cross validations). **M** R^2^ from different univariate models; Regressor angles are ranked by the R^2^. Above the red line are the top 30 regressors, angles from digit IV and V. **N** The top 16 angles. Note that all involve edges from digit V **O** digit V positions corresponding with low and high pain sensitivity calculated from two models in N. n = 34 animals (A-K), n = 58 (L-M), veh = vehicle control, no. inj. = non-injected group

**Figure 5.**
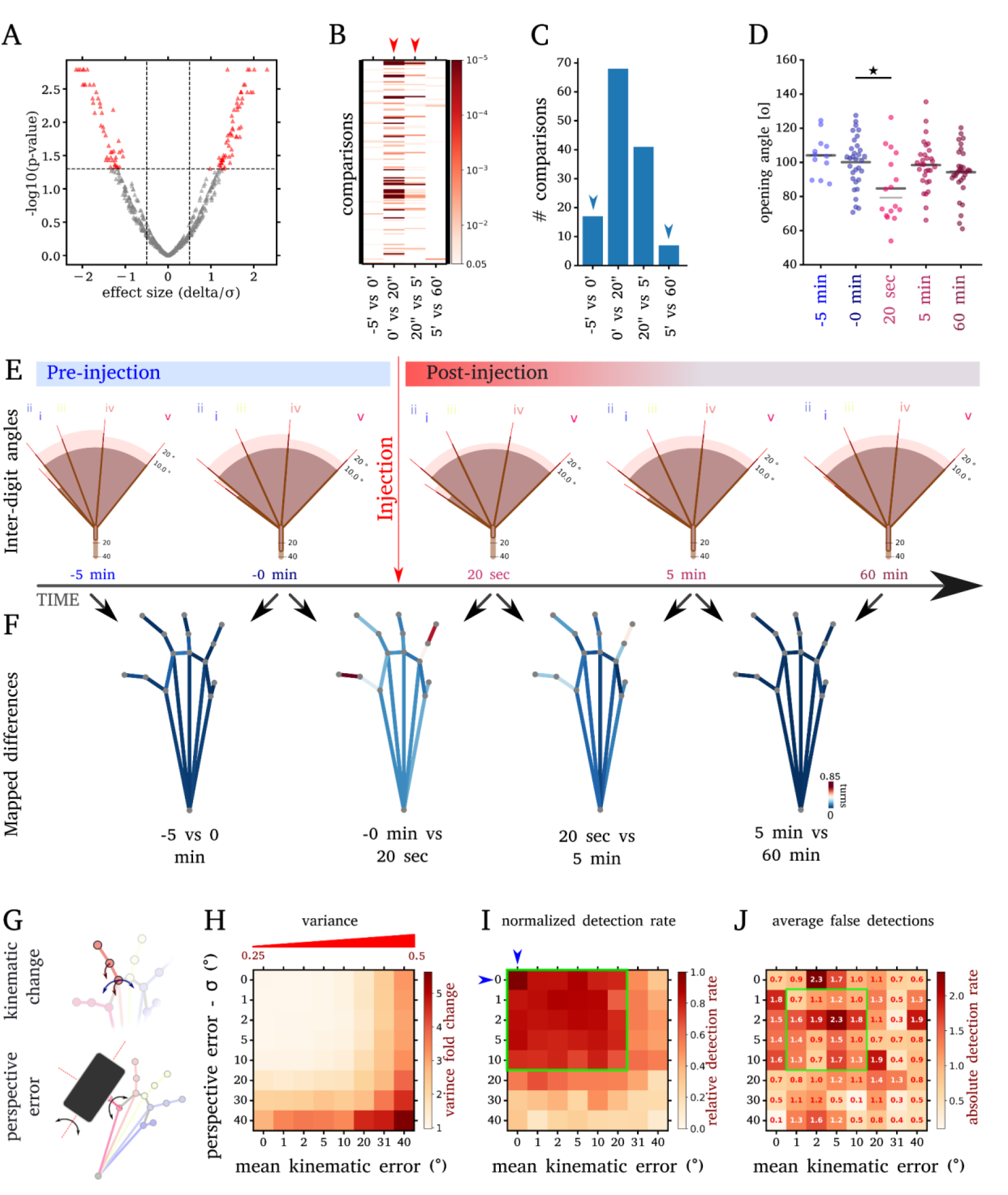
**Habituation and error estimation**. Contralateral paws (control) of injected mice at different time points show significant changes **(A)**. These changes are widely associated with the post-injection time points (**B**, red arrowheads). There are few significant differences between pre-injection timepoints and timepoints occurring well after the injection (**C**, blue arrowheads). **D** In consecutive testing, the TOA significantly deviates only after the injection from the previous group. **(E)** Overall, only moderate change in interdigit angles **(F)** Most of the significant changes occurred following the injection. Consecutive control timepoints show little postural change. **G** shows axes that are varied when calculating kinematic (unrelated movement, blue arrow = pivot angle, red arrow indicates bend angle) and the two-axis perspective error. **H** displays fold change in variance plotted across different levels of perspectives and kinematic errors for two groups of significantly different poses. Note that the fraction of bending segments increases the mean angular kinematic error (red indicator triangle above). **I** Average detection rate across 50 trials. Note the resilience against both errors (green frame. Blue arrow heads denote reference without kinematic or keypoint error.). **J** Average false detections are not systematically increasing due to perspective or kinematic error. Green frame denotes human range of perspective error and moderate task-unrelated movement of the paw. n = 34 animals (A-D), * p<0.05, (permutation tests, FDR corrected)

**Figure 6.**
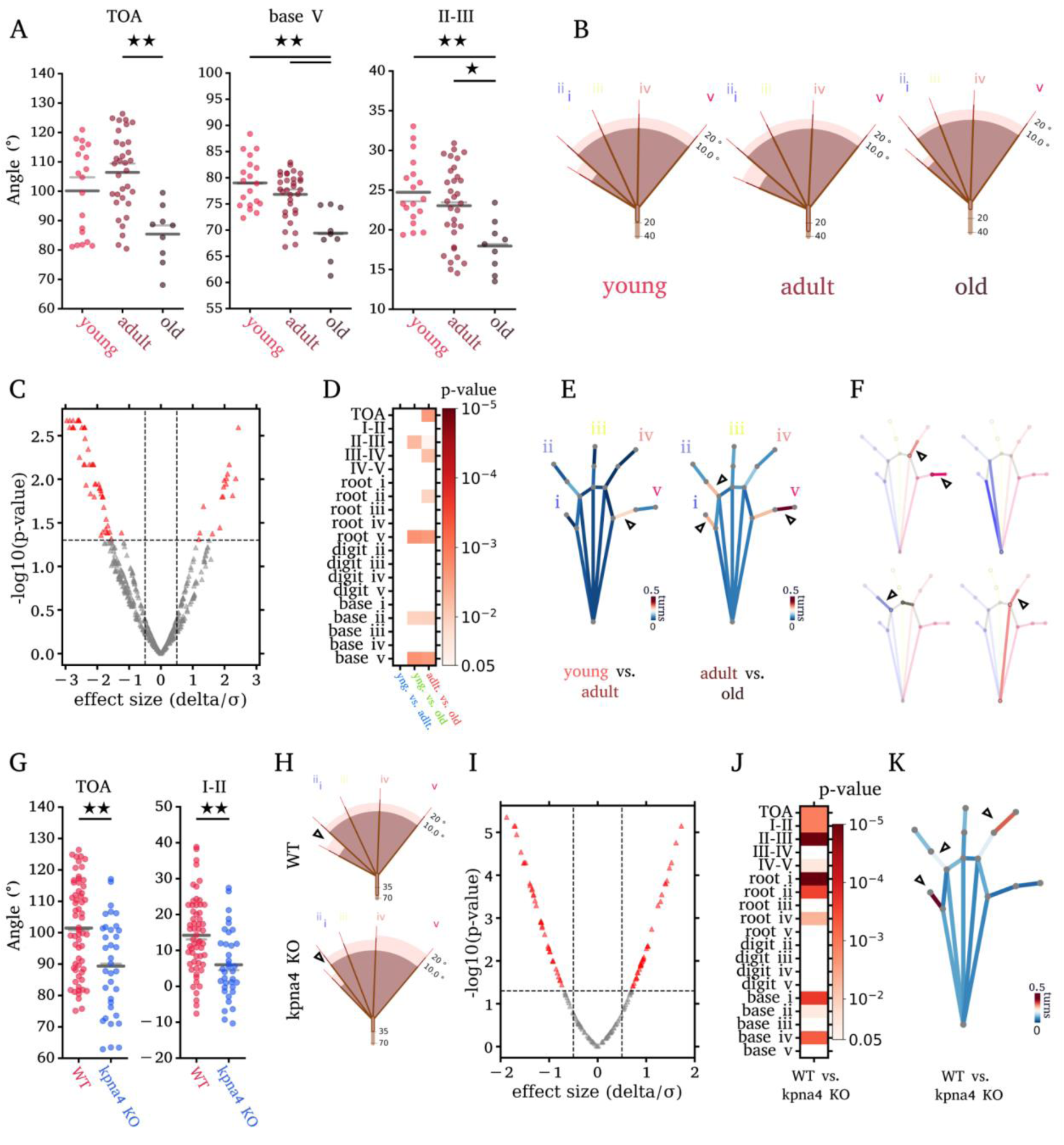
Paws bear a postural aging signature. **A** shows specific opening angles for young, adult, and old mice, revealing a significant decline in paw opening in aged mice (Watson-Williams tests, FDR corrected). **B** inter-digit angles for the age-groups. Comparing all angles (**C,D**) reveals many significant age-dependent changes. **E** Mapping of significant differences onto paw shows changes are strongly driven by digit I, II, and IV, which is consistent with our best-performing regression (**F**) model that uses angles from digits II,IV, and V to infer age (R^2^ = 0.31±0.11), **G** TOA and I-II angle are significantly reduced in kpna4 null animals. **H** inter-digit angles for wt and kpna4 KO animals (note the difference in digit I position, arrowheads). **I,J** Many angles change significantly, much of the significant change (**K**) is associated with digit II, IV, and especially I (arrowheads indicate digits driving postural change). A-F 63 animals (wt), G-K 37 animals (kpna4 KO). * p<0.05, ** p<0.01, *** p<0.001 (Watson Williams test, FDR corrected) young 2-6 months, adult 6-12 months, and old animals 12-20 months.

We next tried to understand if the induced postural change is strong enough to infer the type of treatment from the paw posture. Due to high variability in the first minute after the injection we focused on the 5- and 60-minute time points that we pooled for each experimental group to obtain enough observations for cross-validation. To classify non-injected vs injected paws, we first pooled all the paws that received injections into a single injected group (25 mg, 50 mg, and vehicle control) into one injected group. When subjecting their PC scores (**Fig. 4H**) to a random-forest classification, injected and non-injected paws can be distinguished with good accuracy (**Fig 4I**, av. F1 0.73±0.05). In contrast, when we kept the injected groups apart, RF classification did not accurately discern between the capsaicin-injected and vehicle control groups (**suppl Fig 5C**). This reinforces that the injection itself causes the most distinctive and persistent change. Subsequently, we evaluated whether a regression model could infer capsaicin dosage from the angular geometry, using the vehicle control group as zero and a square transformation of the capsaicin dosage to better reflect the phenotypical similarity between 25 mg.kg^-1^ dosage and vehicle control group. We then screened sine and cosine-transformed angles for the best predictors and found that a univariate model based on single angle from digit III explained ∼56 % of the variance in injected capsaicin dosage (**suppl Fig 5D-G**). Other regression approaches (michaelis menten kinetics or a neural net, **suppl Fig 5J&K**, respectively) did not substantially improve the fit. However, the addition of three more angles (best model, **suppl Fig 5H&I**) increased the explained variance to ∼63 % (**Fig 4J**). Similarly, a multivariate regression on the two best PCs (4,11) explained ∼54 % of the variation in injected capsaicin dosage (**suppl Fig 5L**). These PCs separated the paws receiving 25 mg.kg^-1^ or no capsaicin from the paws injected with the highest concentration (**suppl Fig 5M**), again indicating only minor differences between vehicle control group and 25 mg.kg^-1^ dosage and a more pronounced effect of the highest dosage which is also reflected by the group-specific postures (see **suppl Fig 5N**). Pooling the two posturally similar 25 mg.kg^-1^ and vehicle control groups into a single low-concentration-group and classifying the low- and high-capsaicin-concentration-treated paws based on PCs and 4 and 11 yielded satisfactory performance (0.77±0.7 F1 score, **Fig 4K**). Regression and classification results suggest a posturally large impact of 50 mg.kg^-1^ while the effects of the lower dosage are less distinct from the vehicle control. Overall, the needle penetration alone causes the most substantial change in posture marked by clenching and positional variability over a minute following the injection, while capsaicin further increases this positional variability. Importantly the postural changes allow estimating the administered dosage from the paw posture.

### Digit V posture correlates with von Frey score after SNI

The dose-dependent effect of capsaicin on the paw posture prompted us to ask whether postural features could also reflect pain sensitivity in our SNI dataset. Three months after SNI, virus-treated groups showed recovery from neuropathic pain by exhibiting less pain sensitivity in the von Frey test (**Fig. 1 D&E**). This provided a dataset with varying pain thresholds and paw postures, which we used to determine whether paw postures could explain pain variability. To do so, we regressed log_2_-transformed von Frey scores on different sine and cosine-transformed angles. While searching for the best predictors of von Frey scores, we noted that single angles rather than combinations are better predictors (data not shown). The best predictor could account for 60±0.09% of the variation in von Frey scores (**Fig. 4L, suppl. Fig. 5 O**). Remarkably, among the 30 most predictive angles (**Fig. 4M**, marked by the red line, see also **Fig. 4N**), 29 angles were associated with digit V while only one was associated with digit IV. This observation is consistent with the fact that the sural nerve primarily innervates the digit V and IV to a lesser extent and the sural nerve is spared in the SNI surgery (**Fig 1A&B**). Using the models relating digit V segment posture to the von Frey score, we estimated digit V positions corresponding to high and low pain sensitivity (**Fig. 4O**), as the von Frey test is a surrogate measure for pain sensitivity. This aligns well with our previous finding that digit V contributed with the most significant changes to the overall postural change between injured and non-injured groups (**Fig 3 H**). In conclusion, the posture of digit V, which is innervated by the spared surreal nerve, emerges as a sensitive indicator of pain post SNI and may even serve as a simple, paradigm-specific visual biomarker of recovery from neuropathic pain.

### Paw posture varies little over time

The consecutive measurements of the contralateral side, the non-injected paw, allow us to assess postural change over time and the contribution of habituation. Therefore, we compared angular geometry of the contralateral side between consecutive timepoints (**Fig 5 A-C**). While there are differences between groups, most of these changes occurred only following the injection to the contralateral side (from the viewpoint of the control paw) while reverting to baseline at later timepoints (5 and 60 min, **Fig 5 B,C,D**), hinting at transient contralateral effects of noxious stimuli on the paw posture. It is noteworthy that only 5 minutes after receiving a noxious stimulus to the contralateral paw, the control paw reverted to its baseline posture which it maintained for the next hour. The small, transient nature of these changes underscores the minimal influence of stimuli extrinsic to the sensorimotor control of the paw. In general, the effects on the overall paw posture seem limited and are much less profound when compared with pain or injury (compare **Fig 5E** to **suppl Fig 4 & Fig 3E**). Mapping significant geometric changes back onto the paw revealed very little change within the two first time points and between the 5- and 60-minutes time points (**Fig 5C,F**). Little postural change between the first two timepoints after placement in a new environment suggest robustness to environmental change. Yet, the overall lower number of changes between 5 and 60 min as compared to the initial interval argue for some habituation effects. Together, these observations indicate that the paw posture is robust against disruptive stimuli unrelated to sensorimotor control of the paw by exhibiting transient low amplitude change that rapidly stabilizes.

### Detection rates are resilient

Observing significant changes in consecutive postures, we wondered how many of these arise from random alterations in the virtual skeleton. Such changes might stem from task-unrelated digit movements, perspective errors when acquiring the images, and keypoint placement errors of our model. To address this, we implemented a kinematic chain transformer that applied random digit segment movements to an increasing fraction of digits segments (0.25-0.5) with dependent segments co-transformed and progressively larger bend and pivot angles (**Fig 5G**). The resulting 3D skeletons were projected on a 2D image plane with added random pitch and roll errors modelling camera alignment errors (**Fig 5G**) We further perturbed the skeletons by adding the corrected keypoint placement error of our model. This allowed us to systematically vary digit movement, perspective error, and keypoint wobble to test how those would affect our ability to detect significant angular change in paws. We then introduced moderate to large-scale changes to a single virtual skeleton modelling the underlying differences between two experimental groups. The transformed and non-transformed versions of the virtual skeleton served as the seeds to generate two distinct sets of ten paws each representing posturally divergent experimental groups. Each set was further varied by the kinematic transformation chain, perspective, and keypoint error and then subjected to hypothesis testing as described above. This design allows us to ask how many of the significant differences that are detectable without kinematic and perspective error can be retrieved when error terms are added. As expected, rising kinematic and perspective error increased the combined variance by several-fold (**Fig H**). Nonetheless, the detection rate was robust, so that 10 degree of deviation allowed detecting ∼80 % of the significant changes for both perspective and kinematic error (**Fig 5I**, green frame). To determine the false detection rate, a single skeleton was used to seed two groups with no postural differences. Without error terms added (only keypoint wobble), we detect on average 0.7 angles as false positives. Within the range of human alignment error of a few degrees^23–25^ and a wide range of kinematic error the average false discovery rate is 1.39±0.74 (for n=10), representing a practical cut-off value for this method (**Fig 5J**, green frame). Although the error terms moderately increased the false detection rate, the increase was not correlated with the error terms themselves and therefore did not lead to a systematic rise in false positives (**Fig 5J)**. This indicates that increasing random perspective errors and kinematic errors promote the false negative rather than the false positive rate. Taken together these results suggest that while angular change detection is overall sensitive to perspective and angular error, it is also resilient to it within the human range of accuracy and to a large degree of task-unrelated movement.

### Aging changes paw posture

Aging leads to sensorimotor deficits in the elderly^26,27^. We wondered if age-dependent changes in paw posture could be detected in mice. To address this question, we took paw images from animals of different ages and grouped them into young (2-6months), adults (6-12 months), and old animals (12-20 months). As there was little or no overall difference between left and right paws (**suppl Fig. 6**), or female and male paws (**suppl Fig. 7A-C**), respectively. Therefore, we pooled male and female paws and averaged left and right paws. When comparing paws of differently aged mice, there is a significant decline in TOA (**Fig. 5A&B**). Many angles were significantly changed between differently aged animals (**Fig. 5C&D & suppl Fig 7D**). Projecting the significant changes back onto the contributing edges shows that the aging signature is mostly carried by digit I, II, and V (**Fig 5E**), with digit V contributing disproportionally. Although classification of age groups was unsuccessful (data not shown), combining angles from digits II, IV, and V to explain age, yielded a multivariate model account for only ∼30% of the variance (**suppl Fig 7E&F**). While not sufficiently predictive, the contributing angles align well with the significantly altered edges (compare **Fig. 5E & F**), defining an aging signature characterized by a more compact paw posture and pronounced modulation of digits II, IV, and most notably V.

### Kpna4 genotype affects paw posture

As our dataset included postures from paws of kpna4 KO animals and kpna4 is modulator of sensory and motor neurons^15^, we tried to quantify the impact of the kpna4 genotype on the paw posture. Prominently, a significant reduction of the TOA and the digit I-II angle is evident (p = 0.0052 and p = 0.0075, respectively, Watson Williams tests, FDR corrected, **Fig 5G&H**). Although many angles changed significantly (**Fig 5I&J**), the positioning of digits II, IV, and most prominently I seemed to have the most significant impact. When further subgrouping animals into age groups (**suppl. Fig. 8**), we mostly detected differences between adult animals (**suppl. Fig. 8A-C**). However, the TOA across age groups in kpna4 KO animals is similar to old wild type animals and might be reminiscent of early degeneration. These observations suggest that the kpna4 KO alters paw posture mostly through digits I, II and IV and can be read-out using paw posture analysis.

## Discussion

### Keypoint segmentation is a powerful tool to analyze paw postures

In this study, we re-trained a model from Meta’s Detectron 2 to infer virtual paw skeletons from static images, enabling detection of postural differences. Trained on rare injury-related and spontaneous poses from C57BL/6 mice, the model achieved high accuracy and was optimized for generalization through data augmentation. The postural quantification from keypoints is inherently based on 2D projections of 3D objects, making it sensitive to angular deviations of the camera plane during image acquisition. Such perspective errors distort the projections that are the basis of our analysis. While showing that our approach is resilient to alignment errors within the range of human error (**Fig 5H-J**), acquiring aligned images that minimise the perspective error is still a crucial step in the experimental procedure, determining the overall accuracy of our approach. To prevent a compound error of automatic keypoint wobble and perspective error, we implemented a semi-automatic approach that allows for post-hoc correction of keypoint placement. We embedded this correction tool in an UI that facilitates the efficient processing of large amounts of images. The weights of our model and code are publicly available, thereby allowing any potential user to refine the model’s performance on custom datasets, extend our model to other species or integrate it into custom analysis pipelines. Together, the model and UI tool provide the basis for quantitative analysis of paw postures and eliminates the need for manual keypoint placement enabling high-throughput processing of postural data.

### The toe spreading reflex is robust

The spontaneous paw posture is based on a protective reflex. This toe-spreading-response occurs when animals are suspended by their tails^28^ or kept in a neckhold without support. The reflex is mediated by spinal circuits^29^. Due to this spinal sensorimotor control rather than cortical control, behavioral variability is reduced, and habituation is unnecessary. Under these conditions motor circuit effects dominate and become behaviorally observable, enabling us to quantify the spontaneous postures, injury, and even acute pain with this simple assay. In contrast, goal-driven toe movement (such as foot placement and grip adjustment) requires cortical inputs^30^. Such tasks are traditionally studied using the catwalk paradigm^13^ but they can also be studied by postural segmentation of the paw. This would require more experimental effort because of the higher complexity in behavioral control. The robustness conferred by the reflex is evident when observing unstimulated control paws where repeated measurements after placement in the testing environment showed only minimal postural change (**Fig 5A-F**). Although some postural variation occurred, paw postures were largely maintained despite the novel environment and transient disturbance by a painful stimulus to the contralateral paw, indicating strong resilience against external stimulus. While a dedicated habituation phase is beneficial, the inherent robustness allows reliable measurement during routine animal care procedures with minimal or no habituation. These observations highlight that the reflexive nature of the protective toe spreading makes it a robust model to study the effects of hindlimb spinal sensorimotor circuits by paw posture estimation.

### Paw posture indicates sensorimotor deficits post injury

Quantifying motor and sensory function is reliant upon well-calibrated assays that measure behavioural responses that underlie the complex control. While these assays deliver precise information on clearly defined sensory and motor processes, they require disciplined handling while complexity and variability of animal behavior still can confound the behavioral metrics^31–33^. In contrast, deep learning-based quantification of unevoked behavior has enabled quantifying sensory and motor performance without the need of the animal actively attending to an experimental task^34^. By applying keypoint segmentation to assess post-injury paw posture, we found that paw posture reliably distinguishes injured from non-injured paws even three months after SNI (**Fig. 3A-C**), revealing a characteristic closed posture in injured paws (**Fig. 3C**). This posture and the associated TOA are so distinct that it can serve as a visual marker for injury confirming our initial intuition (**Fig 1F**, **Fig 3B**) highlighting that paw posture is a robust metric for SNI and sensory-motor deficits. We further showed that viral kpna4 knockdown in sensory and motor neurons before the SNI led to partial recovery and reduction of neuropathic pain^15^ while selective depletion in sensory neurons alone after the injury did not achieve this effect (**Fig 3K&L**) on the TOA. On the one hand, this observation demonstrates partial recovery of motor function due to kpna4 depletion, suggesting kpna4 limits axonal regeneration or neuronal survival post SNI. On the other hand, it indicates that the TOA or postural quantification as a reliable metric for motor function in the context of injury. In conclusion, we demonstrate that quantifying the paw geometry can complement or replace traditional assays. The simplicity of our approach allows for integration with routing animal care thereby significantly lowering the experimental effort, overall cost, and irritation to the animal.

### Paw posture indicates the presence of painful stimuli

Animal posture can reflect discomfort. AI-powered analysis of unevoked behavior^6,7^ and behavioral sequences^8^ have been successfully used to estimate pain. Specifically, hindlimb posture has been shown to reveal neuroethological signatures of pain and analgesia from spontaneous behaviors^8^. Our observations show that even static paw posture can reveal intricate details of the animal’s noxious experience. Hind paws respond to needle penetration by transient clenching (**Fig 4A&B**); needle penetration leads to a more robust postural response while capsaicin presence further exacerbates injection-specific postures (**Fig 4D-G**). The capsaicin-dependent postural change was so pronounced that the capsaicin dosage can partially be inferred from a series of angles based on the lower segments of digit II and III and their connections to the heel suggesting those angles as a read-out for capsaicin-induced noxious sensations. It is important to note that neither classification nor regression distinguished postural differences between paws receiving vehicle control injections and those treated with the lower capsaicin dosage well whereas the 50 mg.kg^-1^ dosage was posturally clearly distinct. The 25 mg.kg^-1^ dosage may not have produced a sufficiently large additive effect beyond the injection itself to evoke a differentiable pain stimulus or a subsequent postural change. Pain, an internal and unobservable variable, may exhibit non-linear dynamics^35,36^. Together with the behavioral variability such non-linearity may have obscured differences in low-dose range. Nonetheless, low and high capsaicin dosages were analytically distinguishable, strongly supporting paw posture as a read out for painful stimuli such as capsaicin injections. Moreover, the amount of capsaicin injected correlates with the resulting pain sensation^37^, supporting hind paw postures as a suitable proxy for acute pain sensation. Similarly, when correlating post SNI von Frey scores, a measure for pain sensitivity, with angles descriptive of digit V posture, we found a substantial association (**Fig 4L**). In our surgical treatment, we severed all motor fibers and spared the sensory fibers of digit V. Yet, the mobility of digit V three months after injury raises the questions whether motor fibers of digit V persisted in the sural nerve or regenerated. In any case, low pain sensitivity was associated with spreading of digit V (**Fig 4O**) providing a visual biomarker for post SNI pain sensitivity. Taken together, these results identify paw posture as a visual indicator of hind limb pain, detectable both for simple visual inspection and automated quantification. This suggests paw posture as a robust read-out for pain states.

### Age and kpna4 genotype modulate paw posture

Aging is widely associated with changes in posture and movement^38–40^. Consistently, we detect changes to the paw posture when transiting from adult to old animals (**Fig 5**). This aging signature is evident by a reduction of the TOA and positioning of digit I, II, and V, where the reduction in TOA is largely driven by digit V. Altered or reduced motility of paw digits in aged mice is consistent with age-dependent general decline in sensorimotor ability in animals^41,42^. Similarly, when we compared paws from WT and kpna4 KO animals, we found an age-dependent reduction of the TOA (**Fig 5**). Interestingly, when taking age into consideration, all kpna4 groups showed a TOA similar to aged WT mice. While this might be indicative of an early decline in motor function, large parts of the TOA reduction are accounted for by a change in the motility of digit I. However, this also highlights for the first time a general role of kpna4 for functional motor innervation of the hindpaw. These observations show that the paw posture can serve as a sensitive read out for age-related and genetic changes in paw posture control.

In conclusion, we showed that paw posture analysis can delineate nerve injury, provide an estimate for hind-paw-associated pain, and indicate the postural signature of aging in hindpaws. In some cases, the postures were so striking that they can serve as an intuitive biomarker for visual inspection such as the TOA or digit V position in injury and neuropathic pain. Importantly, the underlying virtual skeletons were taken from single images of static postures. As such, they are easy and fast to acquire during routine animal handling without the need of time-intensive behavioral habituation. This image-based analysis has the potential to reduce or replace classical sensorimotor tests and thereby the discomfort to the animals. While our analysis is based on static images, our approach is scalable to behavioral sequences derived from video recordings (i.e. foot placement or gripping tasks), which has an even greater potential to reveal underlying biological drivers of behavioral variation in limb posture. Especially in the context of neurodegenerative disease, keypoint- segmentation-based analysis of hand or paw movement could be a promising tool for the prediction and analysis of disease progression.

## Material and Methods

### Animal care

Importin α3 single gene global KO in the C57BL/6 genetic background and WT littermates were grown at the Max Delbrück Center for Molecular Medicine Berlin (MDC), Germany at the Weizmann (Israel) and at the NICO institute in Italy. Data collection is coming from three different locations. The data shown are generated from wild-type and importin α3 KO male mice, from animals in three age ranges: young (2-6 months), adult (6-12) old (12-18 months). All experimental procedures at the University of Turin were reviewed and approved by the *Comitato Etico per la Sperimentazione Animale* and authorized by the Italian Ministry of Health (authorization number: E669C.N.1VE, project title *“Studio sulla morfologia dei neuroni sensoriali e sulla nocicezione in topo in importin alpha 3 KO”*). Procedures performed at the Weizmann Institute of Science were conducted under IACUC approval #34620317-2 (capsaicin injection) and #00830118-3 (SNI and viral injections). Procedures performed at the MDC were covered by the animal application number G0023/22, issued by the Landesamt für Gesundheit und Soziales (LaGeSo) Berlin. All experiments complied with Italian Legislative Decree 26/2014 and EU Directive 2010/63/EU on the protection of animals used for scientific purposes and adhered to institutional guidelines to minimize discomfort and reduce animal use to the minimum necessary to achieve scientific validity.

### Data acquisition

Photographs of animals were taken using iPhone 11 & 14 Cameras (Apple, Seattle, US) and Canon EOS 500D SLR (Canon Inc, Tokyo, Japan). For image acquisition, animals were held in neckfold grip triggering toe-spreading reflex (see **suppl Fig 9** and supplementary protocol). The plantar side of each paw was photographed multiple times with the paw being parallel to the image plane (minimizing angular errors). The sharpest image with best alignment was selected for analysis by an experimenter blinded to conditions. For compiling the keypoint segmentation training set, the camera angle relative to the paw was varied to collect more angular variability.

### Capsaicin injections

WT C57BL/6J mice (2-3 months old) were injected with 10μL capsaicin (25μg/kg, 50μg/kg, or saline control) in the right-hind paw with a 10μL Hamilton syringe. Capsaicin was dissolved in a 10 % (v.v^-1^) ethanol and 10 % (v.v^-1^) Tween 80, in PBS. The solvent also served as vehicle control. Images of both hind paws were acquired before, during (video recording), and at set time points after injection.

### Preparation of training data

959 images of mouse hind paws were taken during routine animal handling or after behavioural experiments as published in^15^. Three expert users annotated bounding boxes and 15 keypoints denoting paws on these images (**Fig 2A**). To offset the imbalance in the relative frequency of paw postures, homogeneity of backgrounds and fur colours, images were subjected to scaling, rotations, translation, and changes in RGB channel ratio. Moreover, composite images containing between 1 and 3 paws on random backgrounds were created. We increased the relative frequency of paws displaying rare postures in our training set by prioritizing their augmentation. Our training data is available at the <u>open science foundation</u>.

### Retraining and evaluation of the Mask R-CNN model

To train keypoint segmentation, we used a Mask R-CNN model (keypoint_rcnn_R_101_FPN_3x) from Meta’s Detectron 2 suite using PyTroch. The model was trained on 80 % of our annotated data and evaluated using 319K iteration with a batch size of 4 images using Pytorch. Average precision for bounding boxes and keypoints was calculated on an evaluation set that comprised 10 % of the annotated images. To further evaluate the precision in keypoint placement we compared the predicted keypoints to the ground truth keypoints placed by our annotators in the evaluation dataset. To quantify the error in keypoint placement we used the MSE of the distance between predicted and ground truth points. To compare MSE values between images, residuals were normalized by the distance between the heelpoint (keypoint 1) and base of digit III (keypoint 7). The model weights are available at the <u>open science foundation</u> (see data availability).

### Processing and quantitative analysis of angular geometry

Analysis of resulting keypoints was done in python using standard packages (numpy, pandas, scipy, scikit-learn). We implemented a post-hoc correction suite with partial UI support (keypoint correction UI) that is available at **<u>github</u>** (see data availability). Virtual skeletons of paws were inferred, and deviations were user corrected. To eliminate unrelated variance stemming from random positioning of the paw and scaling, we rotated all the virtual skeletons and scaled all the paws so that the edge ranging from the heel point to the onset of digit III aligns with the x-axis. We then scaled the paw so that the length of the edge equals one. All virtual skeletons of left paws were mirrored to eliminate variability stemming from sidedness. For analysis of angular geometry, all 153 pairwise angles between edges were calculated between -pi and pi unless stated otherwise. To detect differences between paws, we subjected all angles to pairwise Watson Williams tests or pairwise permutation tests. p-values were adjusted for multiple comparisons using FDR^43^.

### Principal Component analysis

For dimensionality reduction of paw postures, we used PCA based on the flattened and normalized coordinates of the vertices of the virtual skeleton (scaled and generic left). For PCA all virtual skeletons regardless of condition were pooled to account for the maximal positional variance. For subsequent classification, PC scores were derived from this global PCA rather than running a separate PCA for each classification. This approach yields lower classification accuracy but is less at risk of overfitting and represents positional variability more robustly. Eigenpostures were obtained by multiplying the loadings of individual PCs (*L*) with the column standard deviations (σ) and adding back the columns means (μ):

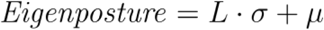

Group-specific postures (*G_g_*) were calculated as the sum of all selected group-specific means of the i-th PC scores 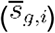 transformed back by the selected i-th Eigenpostures:

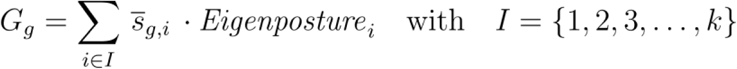

### Classification

We classified paws into injured and non-injured instances based on selected angles using logistic regression. For this analysis angles were calculated between 0 and pi considering only the opening magnitude. Logistic regression was fitted using scikit-learn’s logistic regression module using the L-BFGS-B algorithm for 100 iterations. Model evaluation was done by 30-fold cross-validation. We used random-forest classifiers from the scikit-learn module to classify different behavioral categories based on PCs scores. Between 300 and 800 trees were used during classification. Square root method was used to determine max features (node splitting), trees were grown until all nodes were pure. Generalization error was estimated by the out-of-bag score and average macro F1 score from cross-validation.

### Regression

Angular data was converted by sine and cosine transformation for regression. To mitigate the effects of outliers, we excluded the upper 10 % of data points with the greatest influence on the slope using Cook’s distance. For regression analysis, we used ordinary least squares to fit uni- and multivariate models (scikit-learn, LinearRegression module). The model performance was evaluated by 30-fold cross-validation (50-50 split of data), after fitting the model, the R^2^ was calculated on the held-out portion of the data. We report on the average R^2^ across cross-validation folds. To avoid bias from unequal sample size, the over- represented condition was randomly undersampled to match the number of repeats in the other groups before regression analysis. For multivariate regression models, we selected the optimal combination of regressors (i.e. converted angles or PCs) by a non-exhaustive search. Specifically, we generated random combinations (5-12k) with different numbers of regressors (1-3) followed by selecting the best regressors. Within those top regressors, we performed an exhaustive search over all combinations for models ranging from one to seven regressors, from which we selected the best model.

### Kinematic chain transformation and perspective error

We modeled the kinematic and perspective error by applying a kinematic chain transformation to virtual skeletons. To model progressively increasing but task-unrelated digit movement, 25-50 % of the digit segments were randomly selected and rotated, with dependent edges co-transformed. Rotation angles were sampled from a normal distribution: bend angles (digits curling in) with a mean µ and standard deviation of µ/3, (restricted to 0-90 degrees) and pivot angels (digits pivoting in the sagittal plane) with mean=0 and a standard deviation of σ (restricted from -30 to 30 degrees, also see **Fig 5G**). The resulting 3D skeletons were then projected on a 2D plane, with yaw and roll angles drawn from a normal distribution (µ=0, σ) to model erroneous camera misalignment. To mimic annotation noise, a keypoint placement error (µ=0, σ, normal distribution) estimated from the corrected keypoint placement error (excluding catastrophic placements that would be filtered even by minimal post-hoc correction) was added to the coordinates. Each input skeleton produced 10 error-transformed skeletons, which we then used for hypothesis testing. The detection rate was expressed as the fraction of significant differences preserved under error.

## Author contributions

**CP**: Conceptualization, data curation, methodology, coding, statistical analysis, writing, original draft and reviewing, & editing. **FR**: Methodology, data collection, conducting experiments, reviewing & editing. **GM, SD, PF, NO**: Experimental planning, conducting experiments, methodology, data collection, data curation, writing, reviewing **MB**: writing, reviewing & editing, **LM**: Conceptualization, experimental planning, curation of experimental data, funding acquisition, investigation, methodology, supervision, writing, reviewing, & editing.

## Acknowledgments

We thank Mike Fainzilber for the generous access to his facilities and his support in critically discussing our ideas. We are grateful to Ted Price for the fruitful initial discussion on unevoked pain metrics.

## Data availability

Data, models, and examples are available at the open science foundation: https://osf.io/dc745/

. Code can be obtained from this github repository: https://github.com/ChristianPritz/paw_statistics/

