## Supplemental material for "Paw posture is a robust indicator for injury, pain, and age"

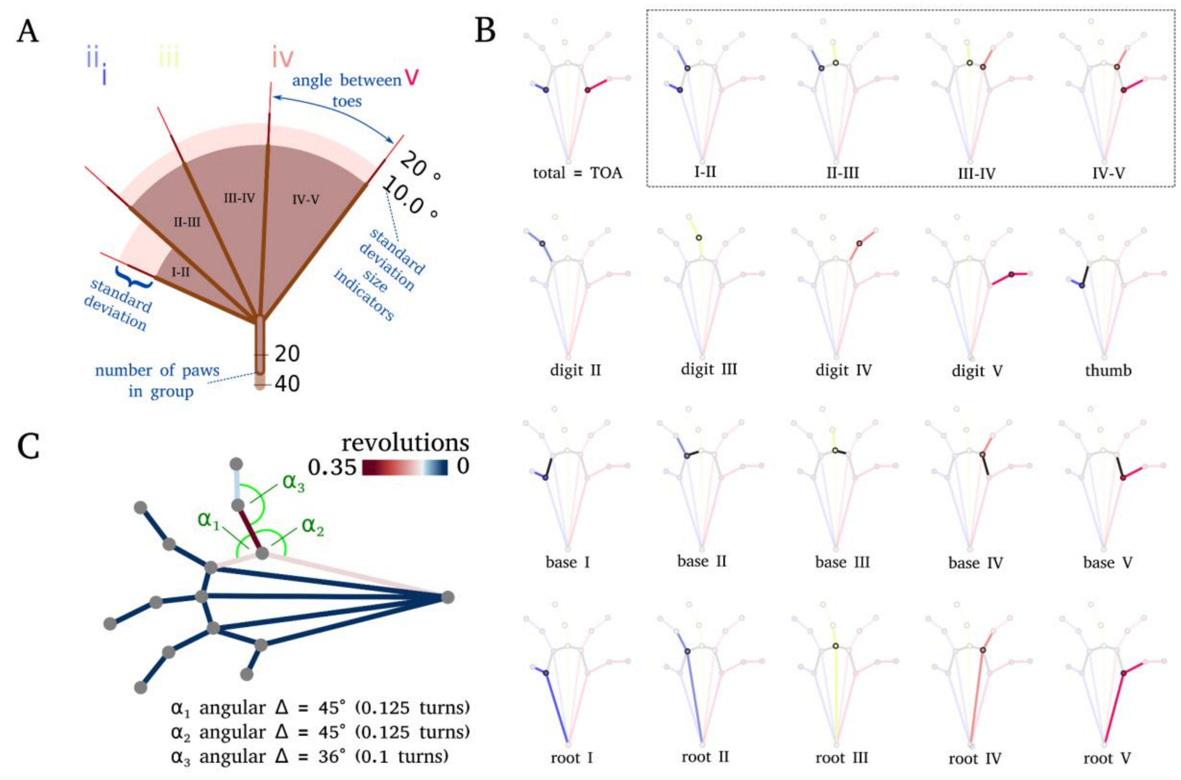


**Suppl. Fig. 1** Metrics used in this study. **A** fan plot of the inter-toe angles of the hind paw. Each slice plot represents an angle between two adjacent toes (see B). The radial extent of the light area indicates the standard deviation of this angle. The stalk of the fan plot denotes the animals included in the group (n). Note that the overall opening angle of the fan plot does not precisely match the TOA because individual toes emanate from different points of origin. **B**) intuitive angles used in this study. Bold lines denote the edges that form the angle. **C** Each angle is defined by two edges, so repositioning a single edge can alter multiple angles. For each edge, the summed angular change of significantly affected angles is color coded. In this example, the edge at the base of digit V contributes to three significantly changed angles (α1–3), while adjacent edges each contribute to one. This summed change highlights the disproportionate impact of the digit V base edge (dark red). Since highly mobile edges can accumulate angular changes exceeding 360°, we report the summed deltas in revolutions (total angular change divided by 360). This provides an intuitive measure of how many full rotations of angular change an edge contributes.

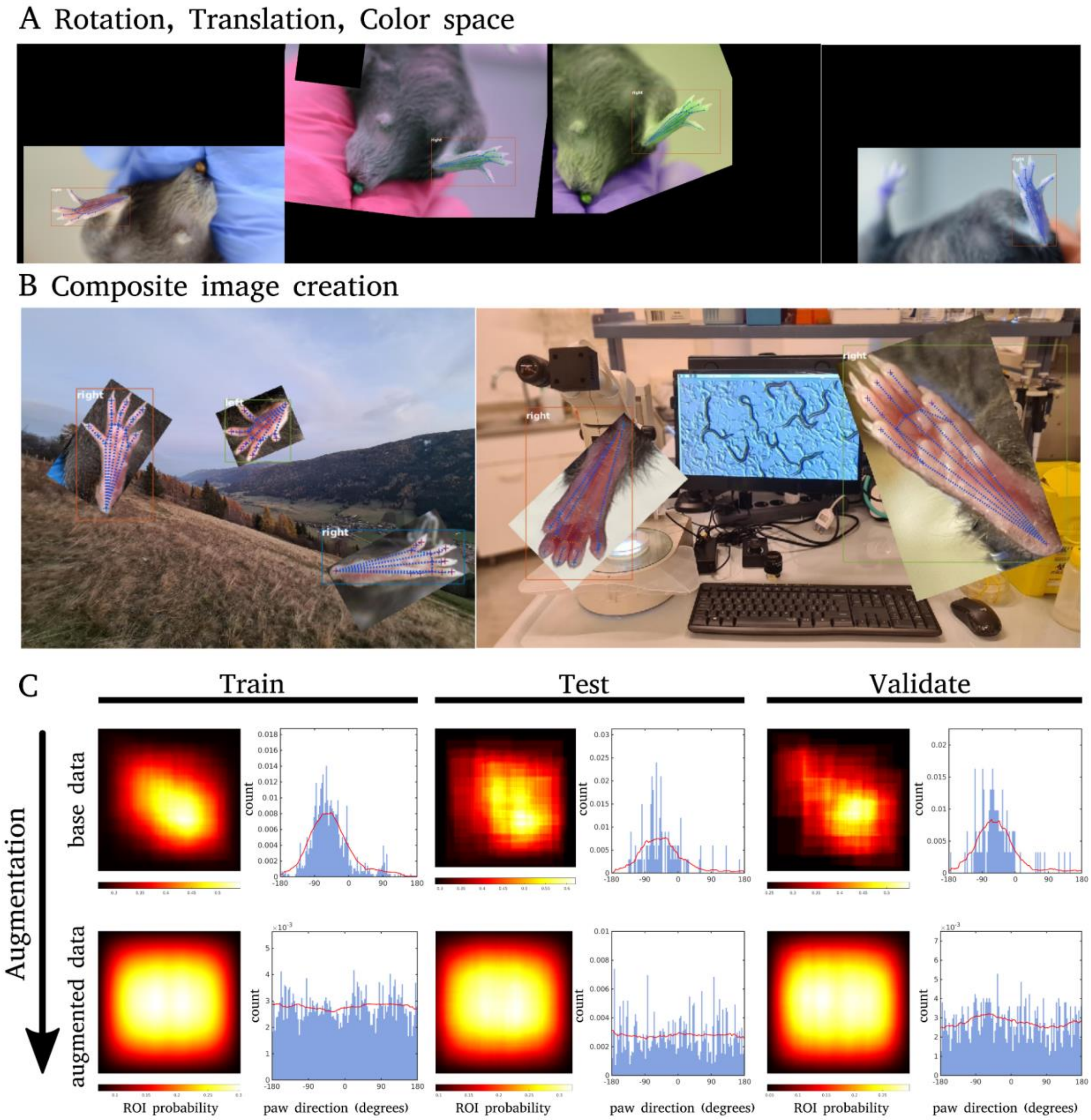


**Suppl. Fig. 2** Augmentation used in our training set. **A** We used rotation, translation, scaling, and color space variation to facilitate generalisation in keypoint placement. **B** To offset homogenous backgrounds in our training set, we created composite images by randomly positioning paw instances in random images. **C** Using amplification, we eliminated the positional and rotational bias in our base dataset. ROI probability denotes the probability of a pixel to be an ROI of an annotated paw. Paw direction denotes that angle of the mid vector (pointing from the heel point the keypoint 7) and the x axis. Heatmaps show roi probability over the normalized image space. Histograms show probability density for paw direction.

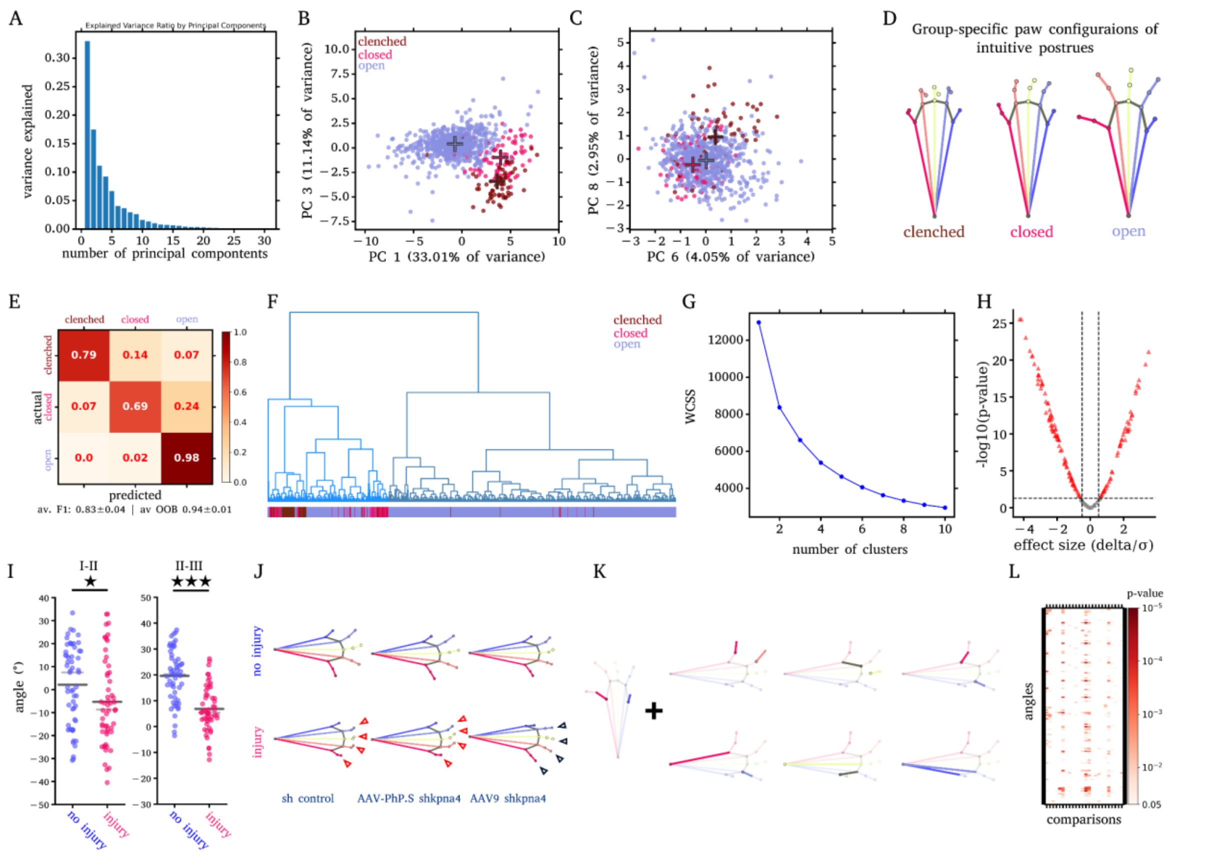


**Suppl. Fig. 3** There are three visually intuitive postures in our dataset. When conducting a PCR on all paws included in this study, 70% of variance is captured by the first 5 PCs (**A**). In some of these PCs (**B,C**) the three postures separate. Calculating the group-specific poses from the centers of these posture groups (crosses in B and C), reflects the stereotypical configuration of these postures (**D**). **E** When classifying these three postures using a random forest classifier based on the PCs (1,3,6,8,10, and 17), we obtain an F1 score of 0.83±0.04 (20-fold cross-validation, n = 871 paws (including time series) from 192 animals). Using hierarchical clustering based on these PC (**F**), we see that most 'clenched' or 'close' paws are clusters. **G** When comparing clustering solutions, we note that no distinct inflection point that would indicate an optimal cluster number consistent with postural variability. **H** Volcano plot of angular comparisons between injured and non-injured paws. **I** Interdigit angles between injured and non-injured paws. **J** shows group-specific postures for the different viral kpna4 knock-downs. Note all uninjured postures display an open configuration (upper row) and that the injured, AAV9-shkna4-treated group exhibits a posture more similar to the no-injury group: straight digits, particularly digit V (black arrow heads). In contrast, sham and AAV-PhP.S-treated groups show a more closed configuration (red arrow heads). **K** Angles that are predictive of injury in logistic regression. **L** Full p-value matrix for comparisons between all treated groups (see **Fig 3J**).

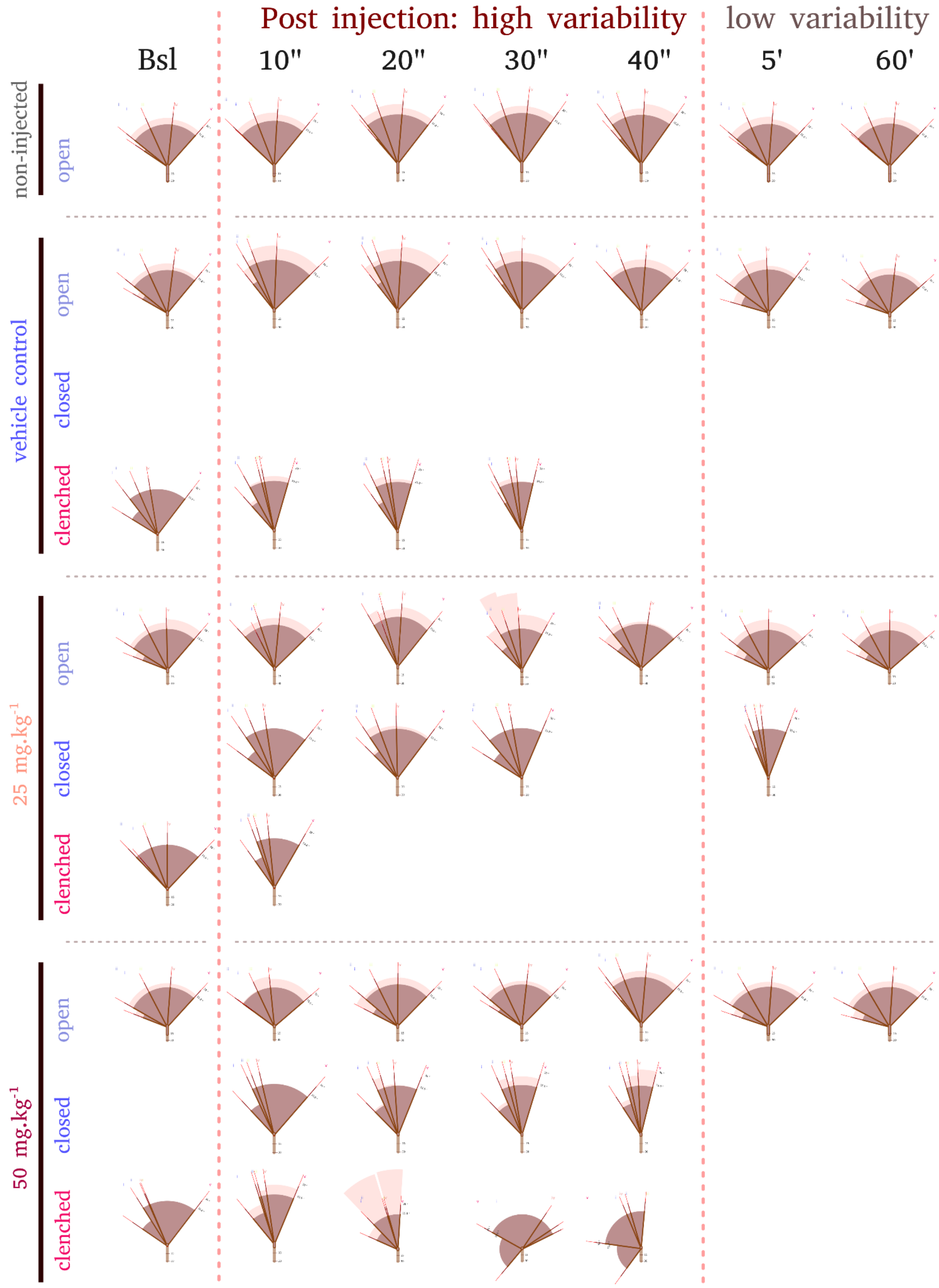


**Suppl. Fig. 4** Angular arrangement of all groups over time (fan plots of inter-digit angles). Note that in the time course of one minute after the injection, there is high positional variability in terms of number of paw postures detected and also inter-digit angles (see standard deviations, indicated by the length of the pink segments). During the 5 and 60 minutes post-injection postural variability decreases. Bsl = baseline level

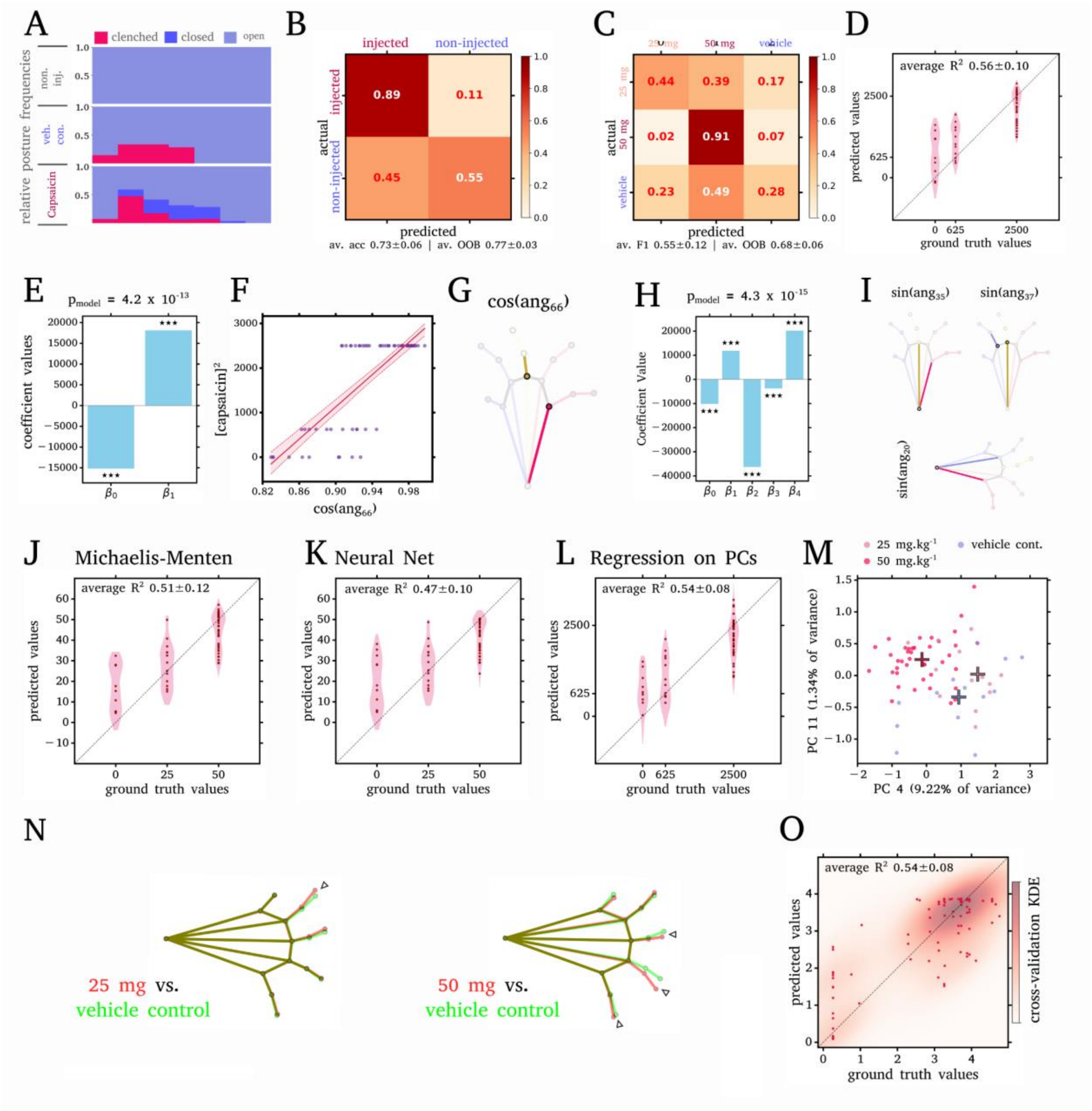


**Suppl. Fig. 5 Acute pain and paw posture: A** Postural distribution after pooling the capsaicin-injected groups. Note that the closed posture does not occur in non-injured or vehicle control groups. **B** RF-classification of injected vs non injected during the first minute after injection. **C** RF-classification of injected groups fails to accurately distinguish injected paws (5 and 60 minutes after injection). **D** predicted vs ground truth values of squared capsaicin concentration for regressing capsaicin dosage against a cosine transformed angle. Note that half of the variance is explained by angle 66 (see G, dots denote predictions from the full model, violins denote predictions from cross-validation trials). **E** coefficients and p-values of the univariate model, **F** data (blue) vs predictions (red line). **G** Edges forming angle 66. **H** coefficients and p-values of the multivariate model, **I** Angles and transformations that together with the angle in G underlie the multivariate model. Fitting Michaelis Menten kinetics

(**J**) and a neural net (**K**, MLPC 1 layer, 20 neurons) to cosine-transformed angle 66 did not improve the fit. **L** Fitting a linear model to PCs 4 and 11 explained ~54% of the variation in capsaicin dosage (predicted vs ground truth, squared capsaicin dosage). These two PCs also differentiate the 50 $mg.kg^{-1}$ from the lower dosage and vehicle group in the PC space (**M**). **N** Overlay of group-specific postures calculated from vehicle control (green) and capsaicin groups (red). Note that the 50 $mg.kg^{-1}$ group deviates more from the vehicle control posture than the 25 $mg.kg^{-1}$ (see arrowheads). **O** Regressing paw angle **145** against von Frey scores from SNI dataset can explain ~60% of the variance in the von Frey score (ground truth vs, predicted values, dots denote predictions from the full model, the heat map prediction density from cross-validation trials).

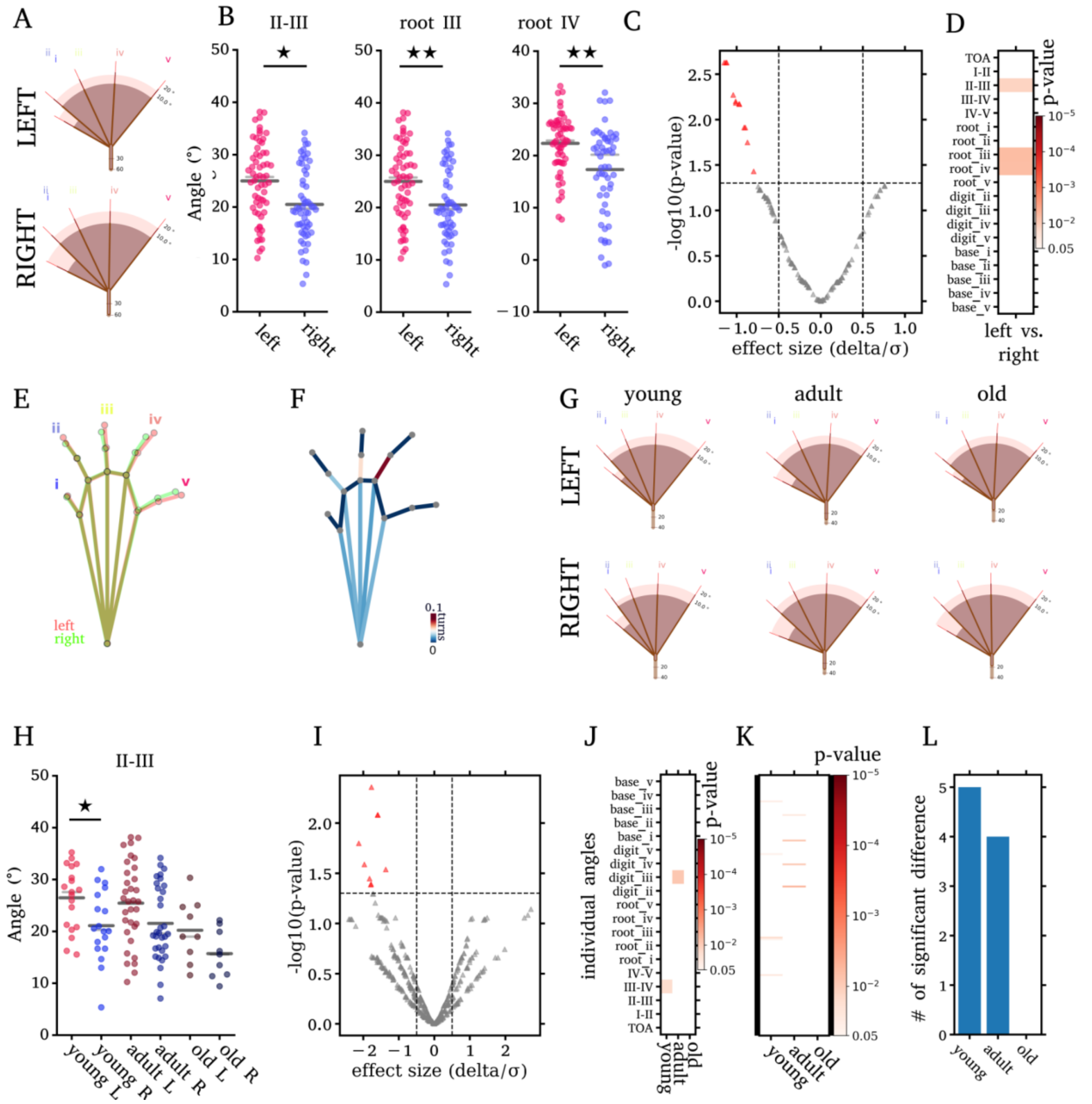


**Suppl. Fig. 6** Differences between left and right paws. **A** Paw fan plot. **B** selected angle scatter plots, **C** Pooling all age groups reveals only few significant differences between left and right paws are marked. Note that while there are significant differences between left and right paws, the number of differences is lower than the number of differences observed when comparing successive timepoints within the non-injected group (see Fig 4 E) **D** p-values from intuitive angles; Strikingly digit III and IV seem to be affected. **E** When overlaying the side-specific poses minor differences are evident. When further differentiating into young, adult, and old animals (**F-G**), even less differences are detectable. These differences are only detectable in young and adults. **F** paw fan plots of all age groups, G Scatter plot of angle II-III, H volcano plot and I heat map of left-right pValues in young, adult and old animals. **K** bar plot of number of significant left-right differences per age group.

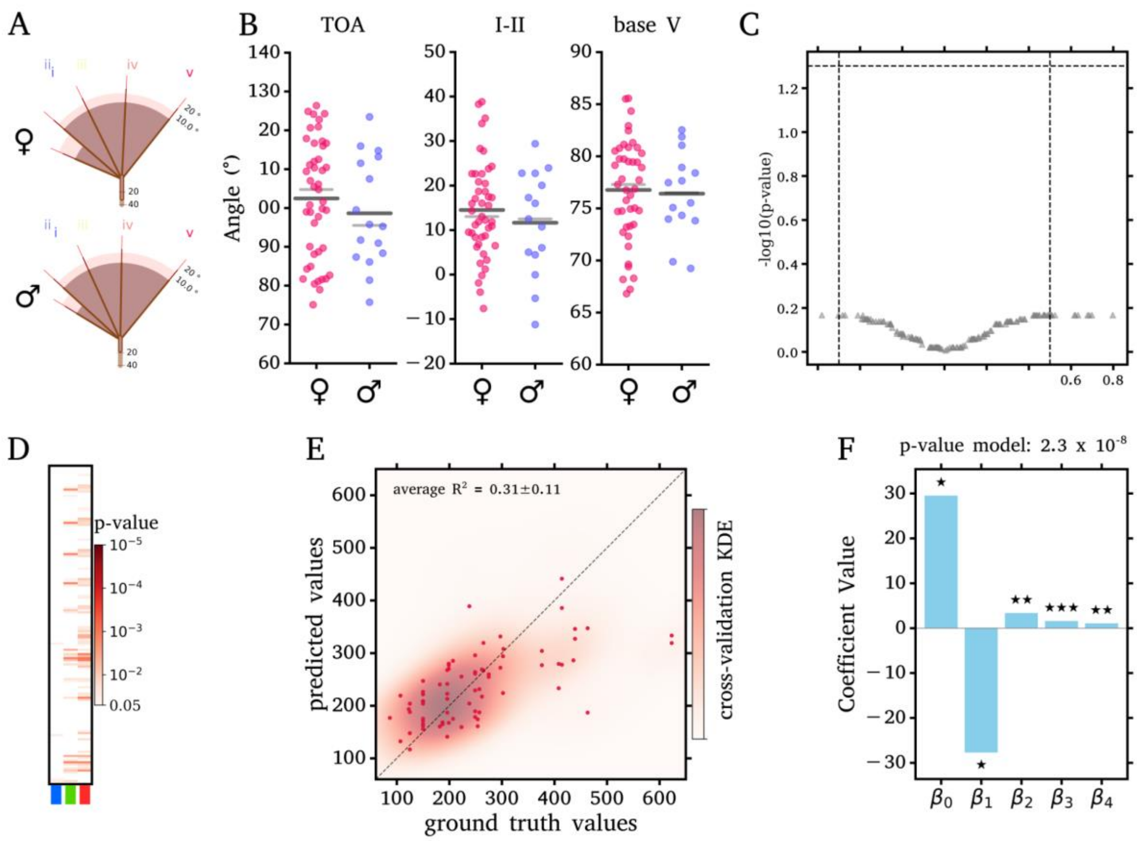


**Suppl. Fig. 7 Comparison between male and female paws.** No striking differences are evident between male and female when considering inter-digit (**A**) angles or representative angles (**B**). No significant differences are observed when considering all 153 pairwise angles (**C**) **D** shows a heatmap of p-values from all angles from between age groups (**see Fig. 5**). **E** prediction vs ground truth values from multivariate regression of angles vs age in days, colored background denotes kernel density estimation over all cross-validation trials. Dots denote deviations of the full model. **F** Coefficients from the multivariate model in E, beta 1-4 are the coefficients for individual angles.

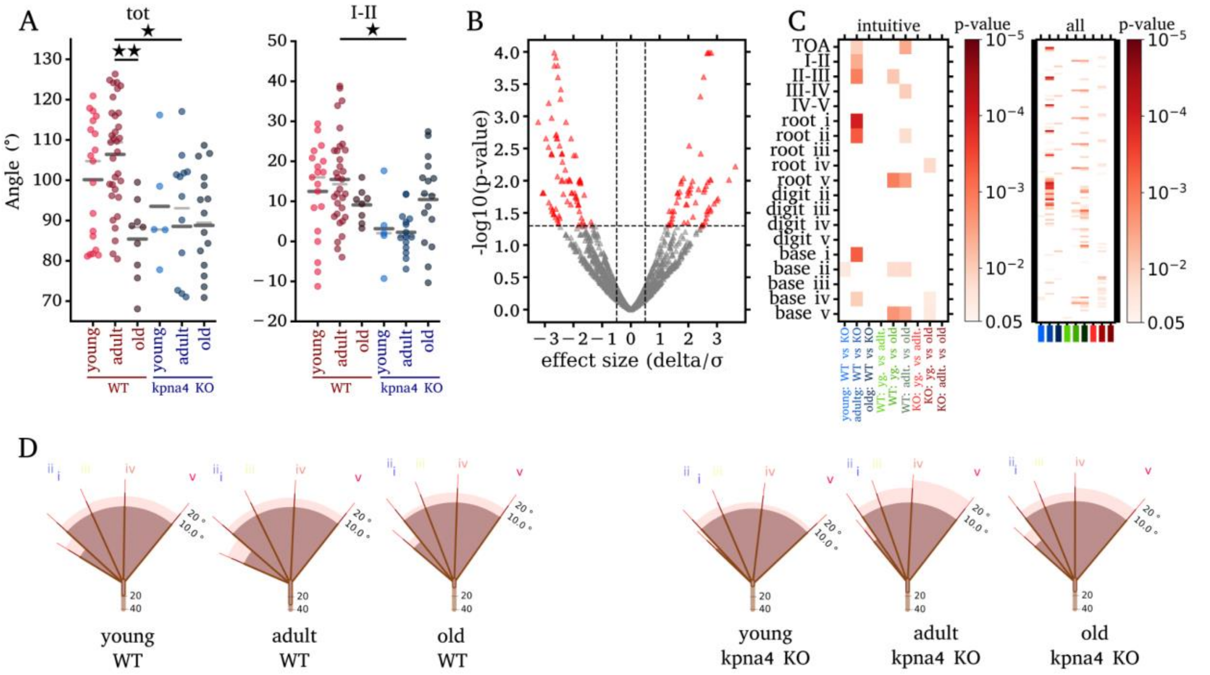


**Suppl. Fig. 8** : Posture differences between wild type and kpna4 -/- animals across all age groups. **A** Scatter plot of selected angles with significant differences. Even when considering different age groups a robust number of angles are significantly different between wt and kpna4 null animals. **B** shows a volcano plot of all comparison **C** displays the p-value matrices for intuitive angles and for all angles. **D** inter-digit angles for all genotypes and age groups.

### Supplemental protocol

**Acquiring images of hind paws for postural analysis**

Paw postures analysis leverages the toe-spread-reflex. The reflex is robust but consider that handling the animal for acquiring images might bias the animal's performance in any follow-up behavioral assay. For imaging, mice need to be positioned so that the reflex is triggered:

- Hold the mouse in the neckfold grip and turn it on its back so that the belly is exposed
- Do not bring any object that could elicit gripping close to the paws. This would strongly confound the result.
- In case the animal is making aversive movements especially with the hind paws, wait until they stop or pick the animal up again.

Note that mice can also be held with the belly facing downwards or held suspended on their tails. While this is probably more efficient in triggering the reflex it makes it harder to acquire images.

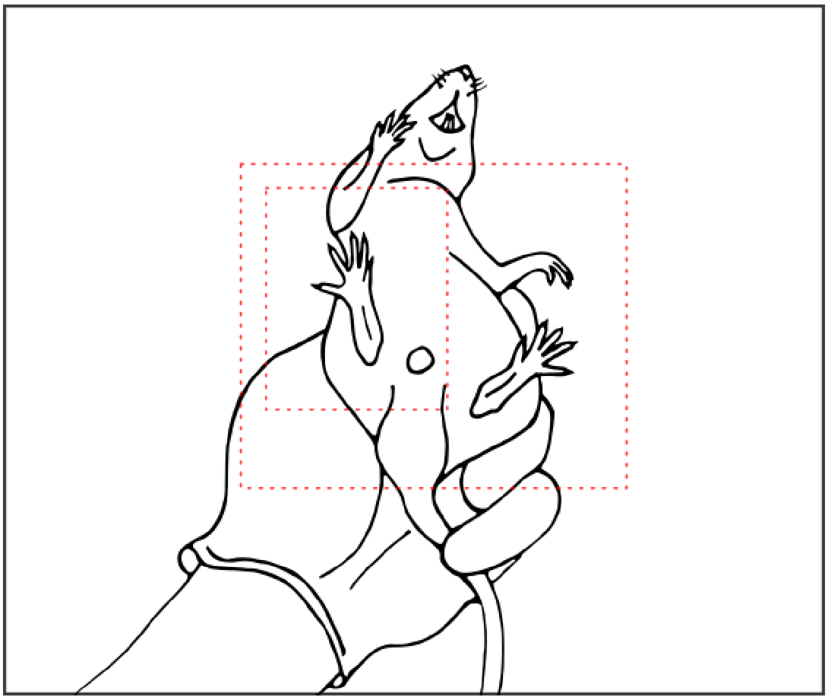

**Supplemental Figure 9:** Handling mice for acquiring images: Belly-up neckhold allows for acquiring images of hind paws. Photographs should show single or both hindpaws close up (red frame denotes optimal image size).

Take an image or record video of the paw. Analysis works best when (**see suppl. Fig 9**):

- The paw is focused
- Image plane and plantar side of the paw are as parallel as possible (this minimizes angular errors)
- The image contains a close-up of one or two paws. If the resolution of the paw is insufficient, the errors in keypoint placement are disproportionately large.
- When taking images it is best to take a series of pictures and select the best shot for analysis.
